## Supplementary Data 1 for "Joint Multi-Ancestry and Admixed GWAS Reveals the Complex Genetics behind Human Cranial Vault Shape"

### Supplementary Information

#### Overview of Genome-Wide Significant Loci

S. Goovaerts et al.

See page 2 for captions

**Overview figures of 30 genome-wide significant loci.** **A**, LocusZoom plots for the segment in which the SNP had its lowest P-value (one-sided chi-square). Points are colored based on linkage disequilibrium ($r^2$) in the 1000 Genomes Phase 3 EUR population. **B,** -log10(P-values) (one-sided chi-square) across hierarchical cranial vault segments. Grey segments did not reach genome-wide significance. **C,** Latent shapes associated with the most significant segment (top row), and full cranial vault (bottom row). Red and blue represent an outwards and inwards deformation respectively relative to the overall average cranial vault shape.

See page 2 for captions

**rs3936018**

**A**

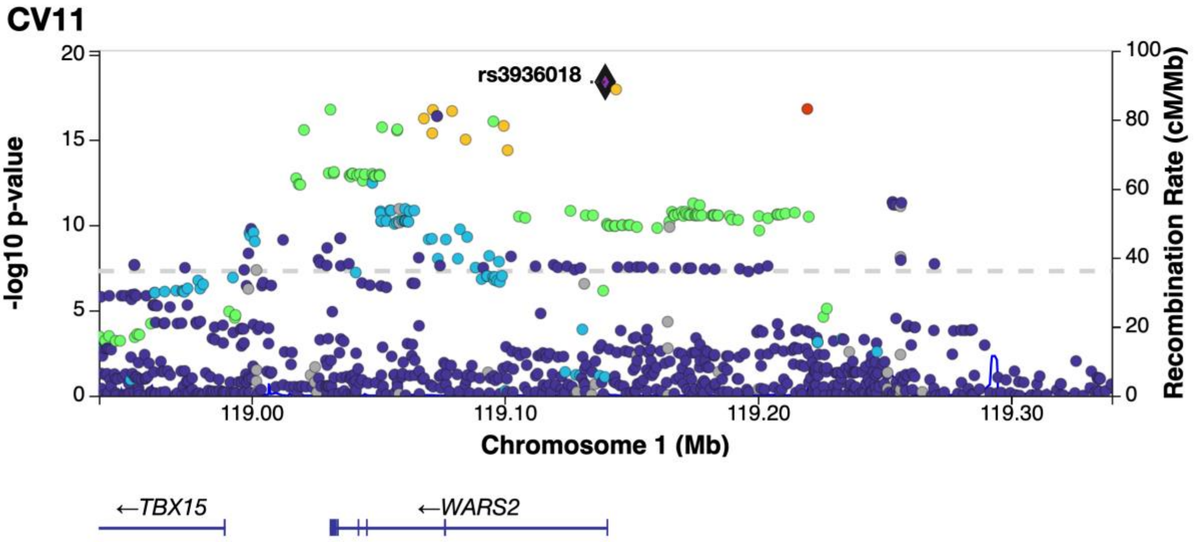

**B**

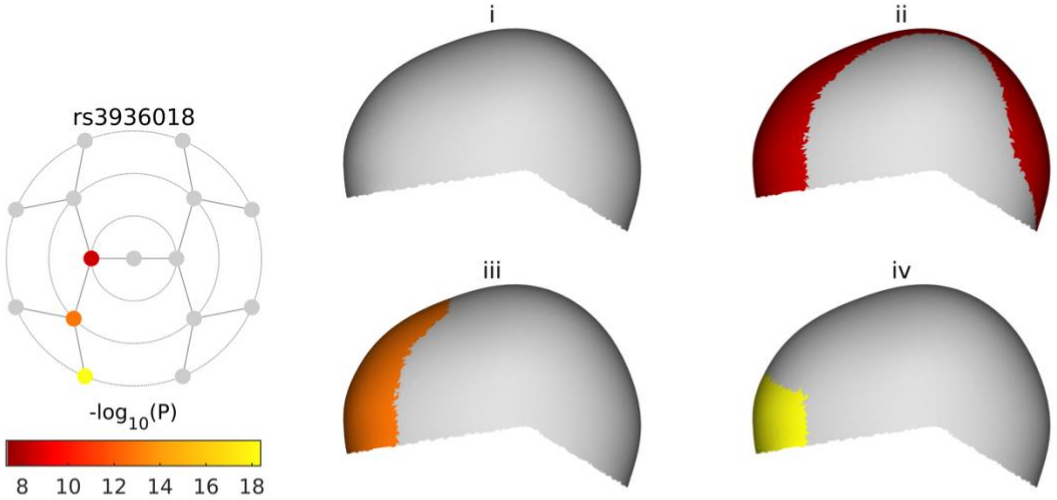

**C**

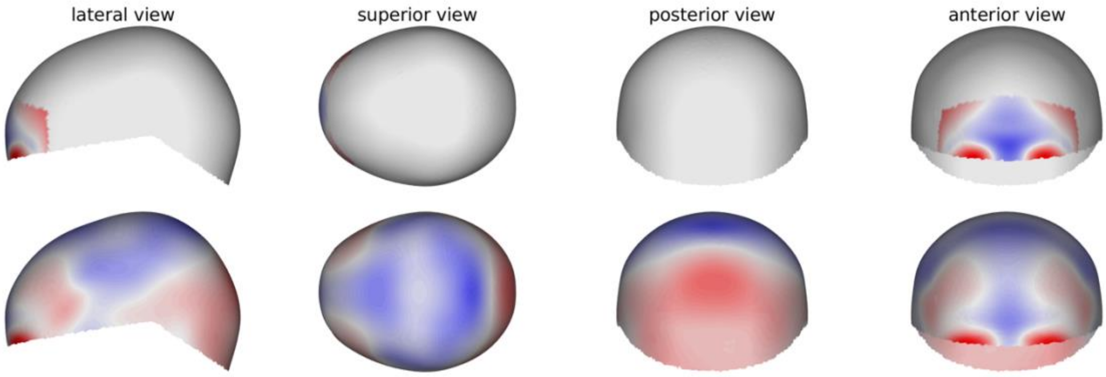

See page 2 for captions

**rs2009778**

**A**

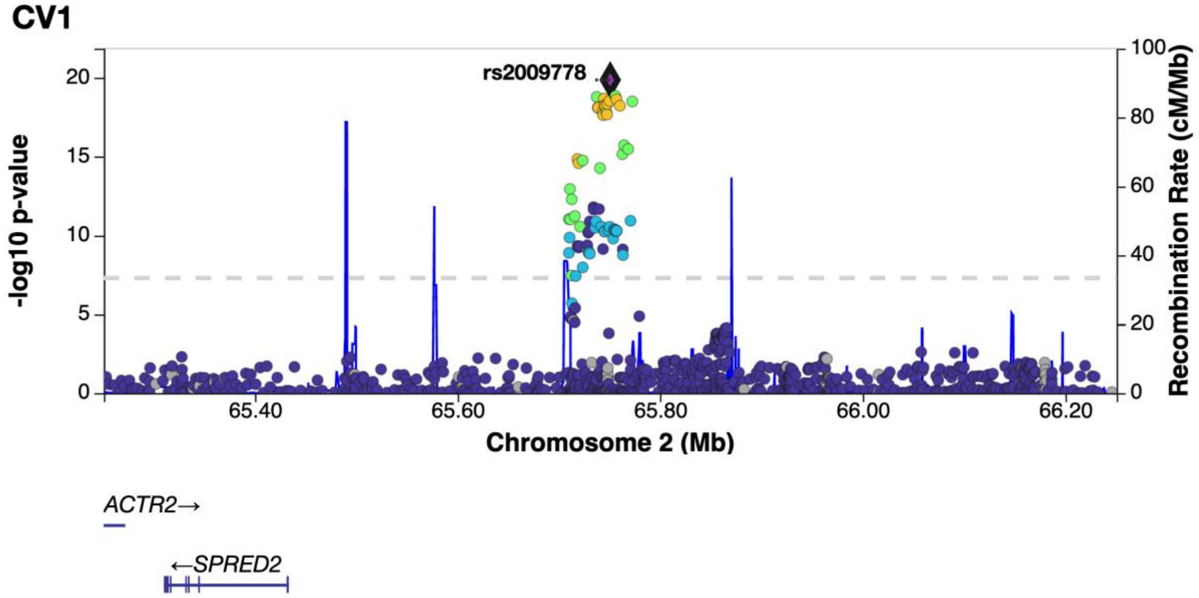

**B**

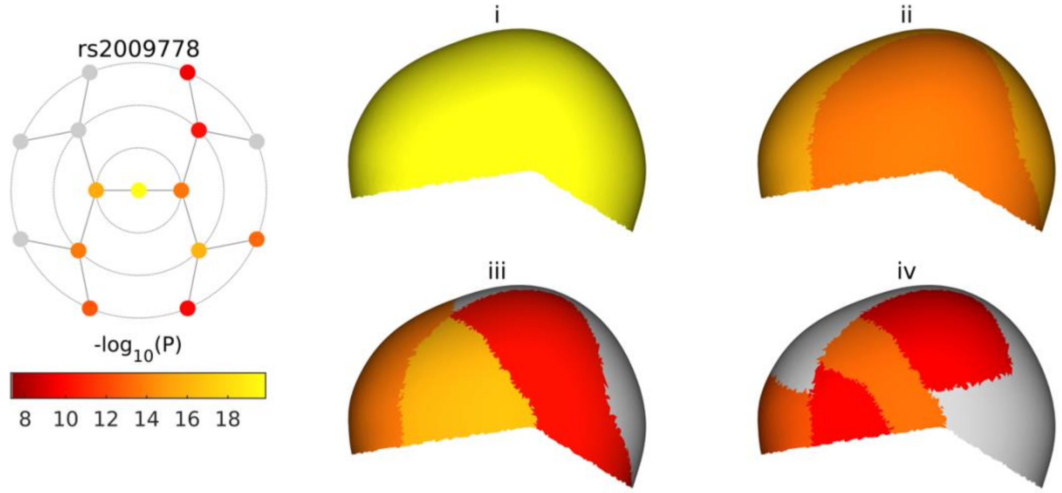

**C**

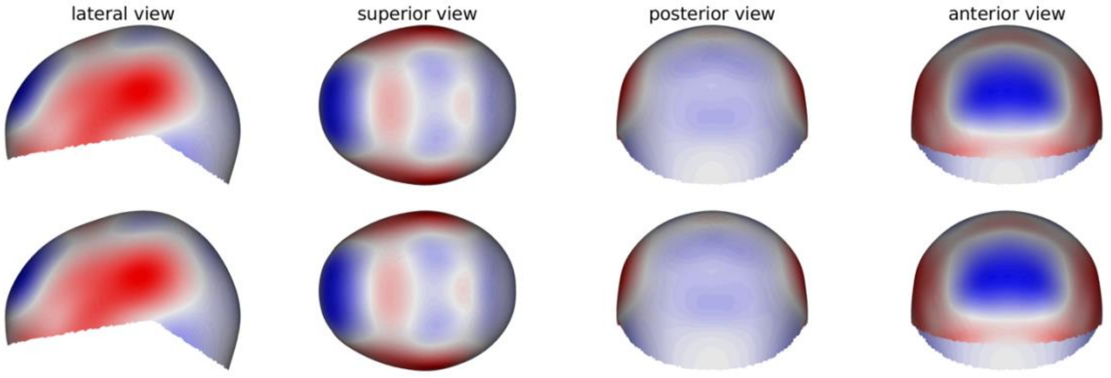

See page 2 for captions

**rs6739488**

**A**

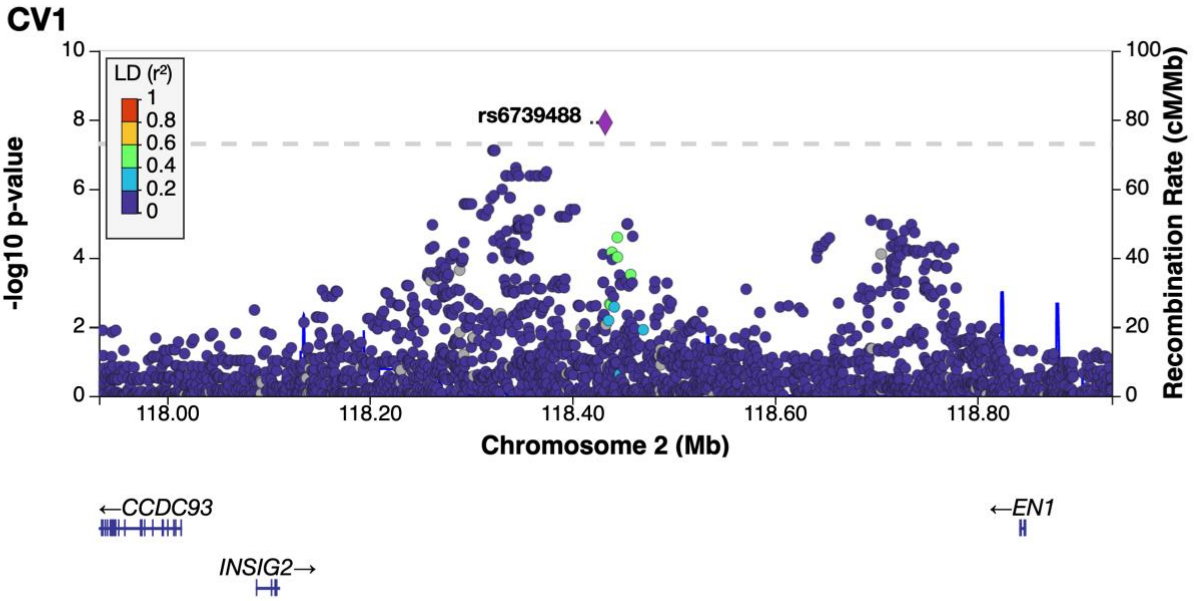

**B**

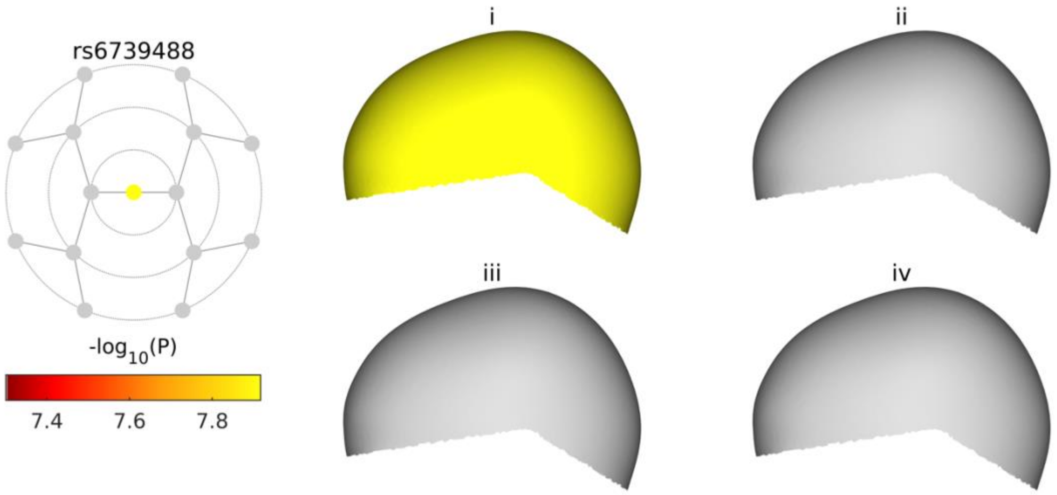

**C**

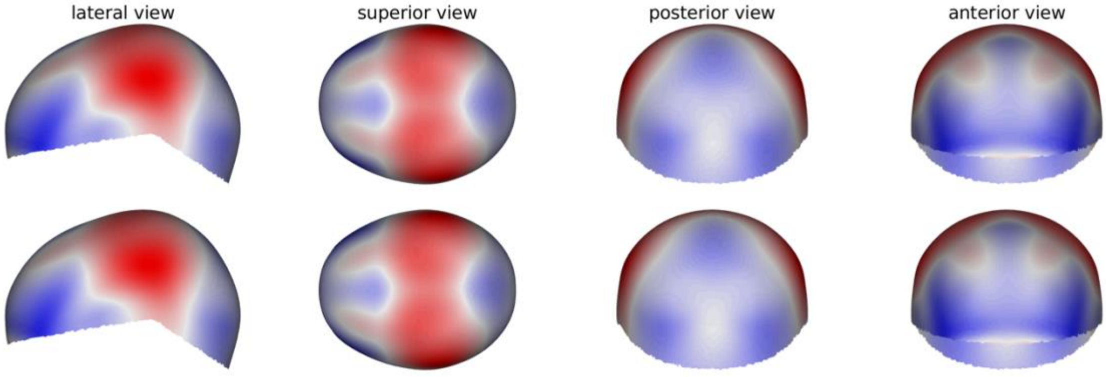

See page 2 for captions

**rs17479393**

**A**

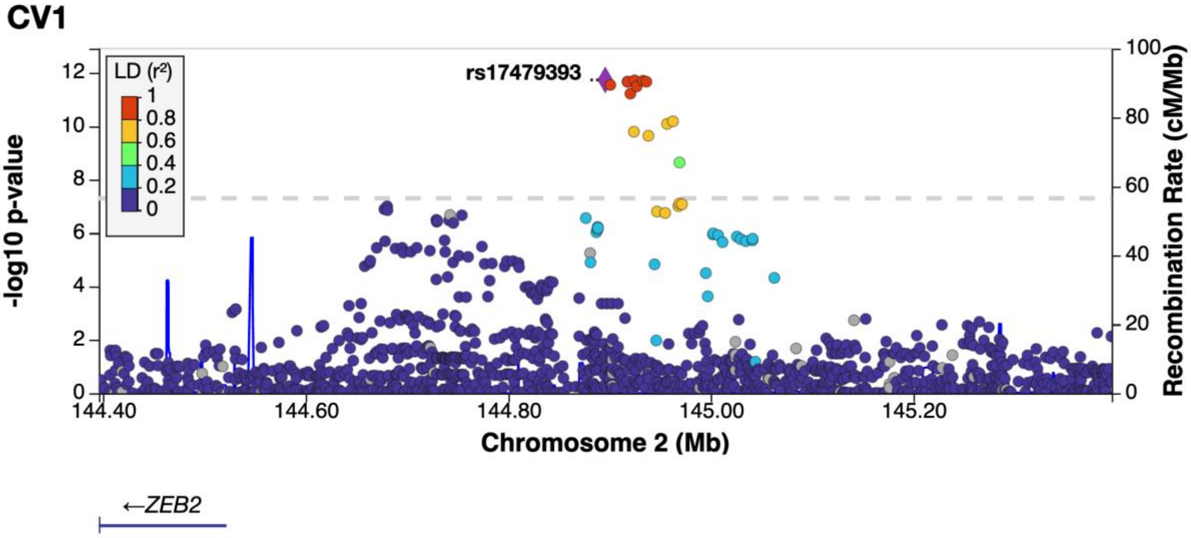

**B**

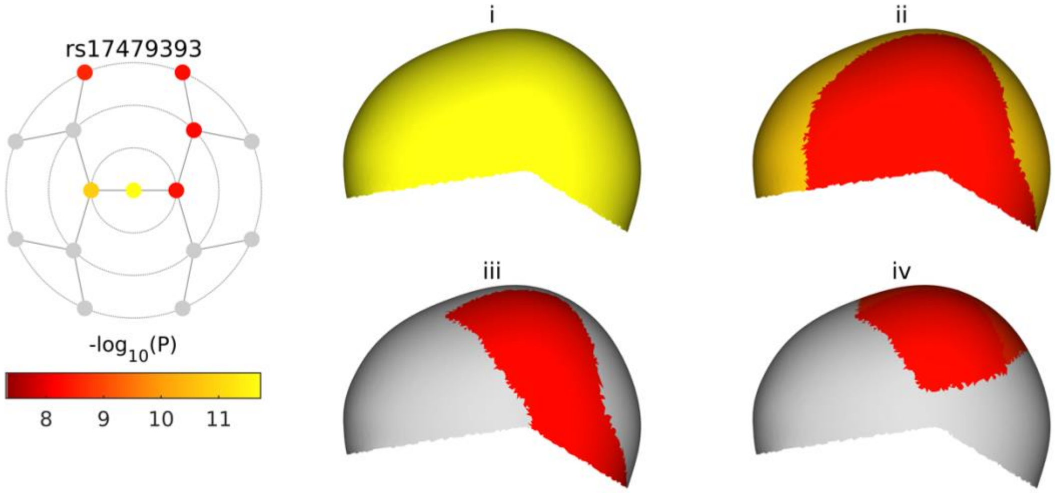

**C**

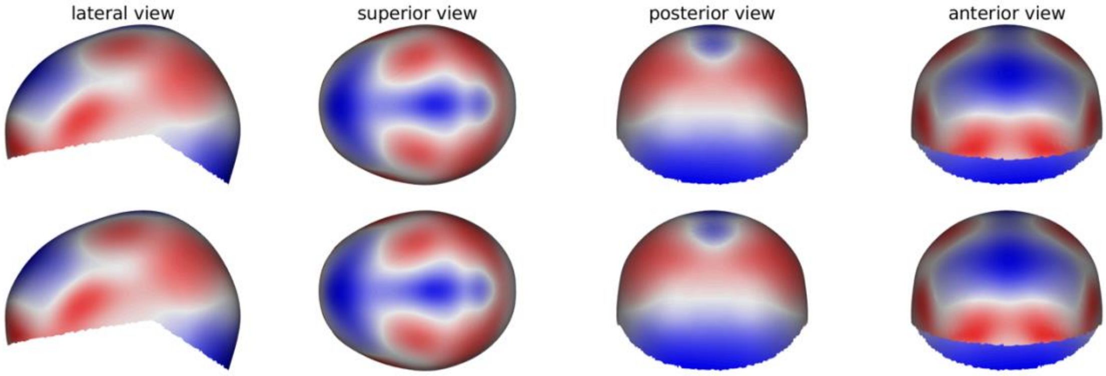

See page 2 for captions

**rs970797**

**A**

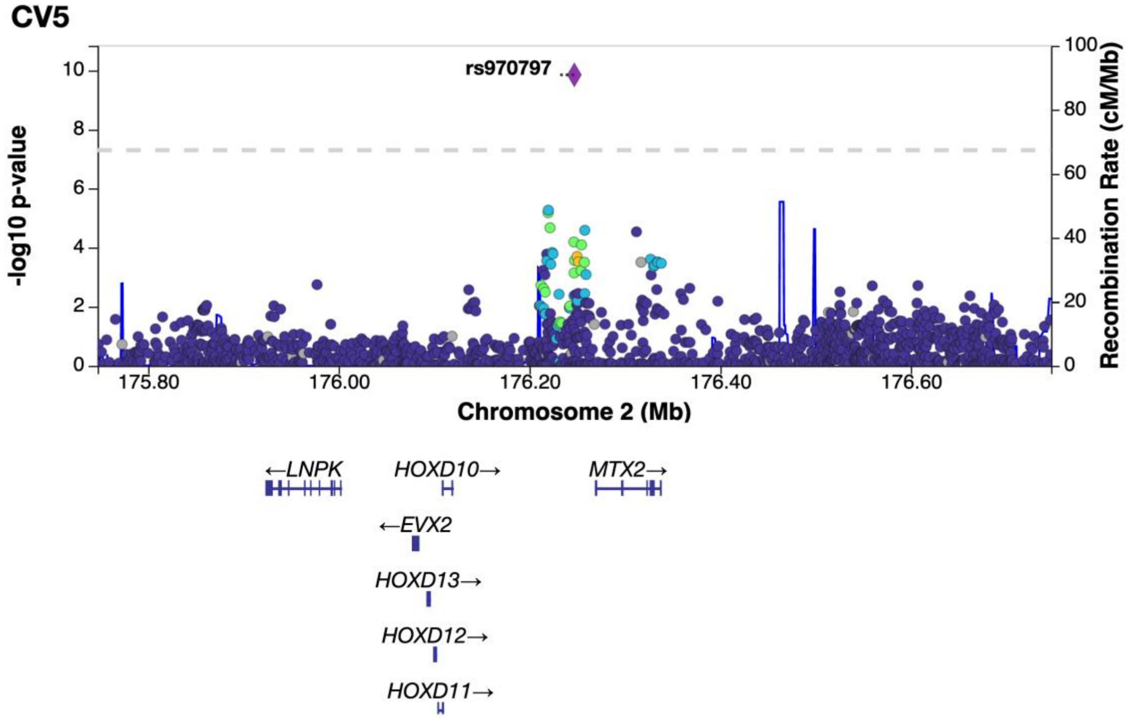

**B**

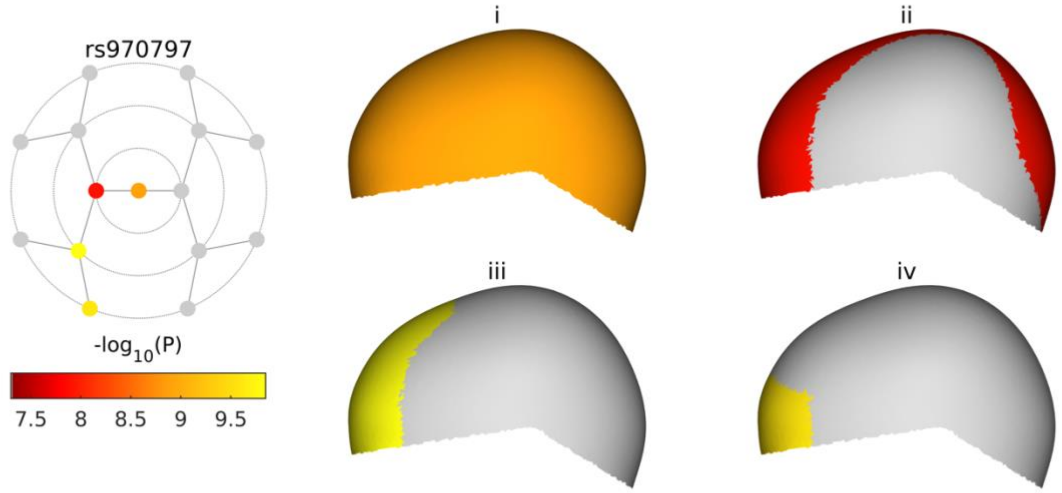

**C**

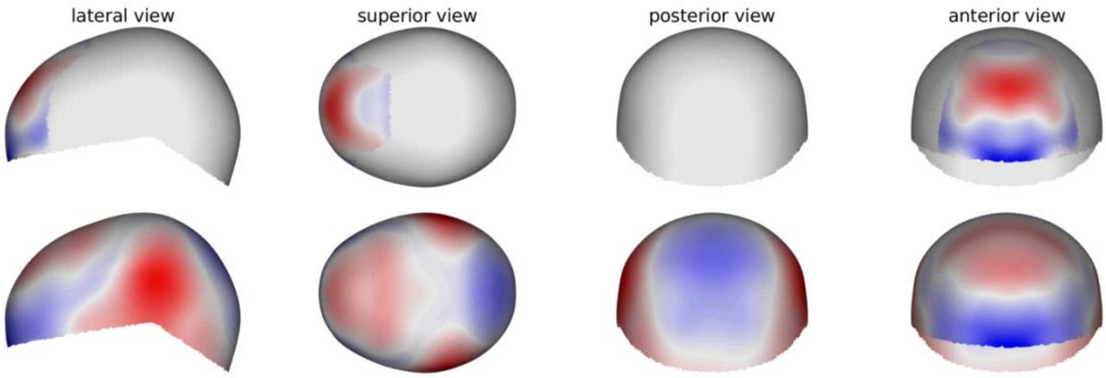

See page 2 for captions

**rs7626244**

**A**

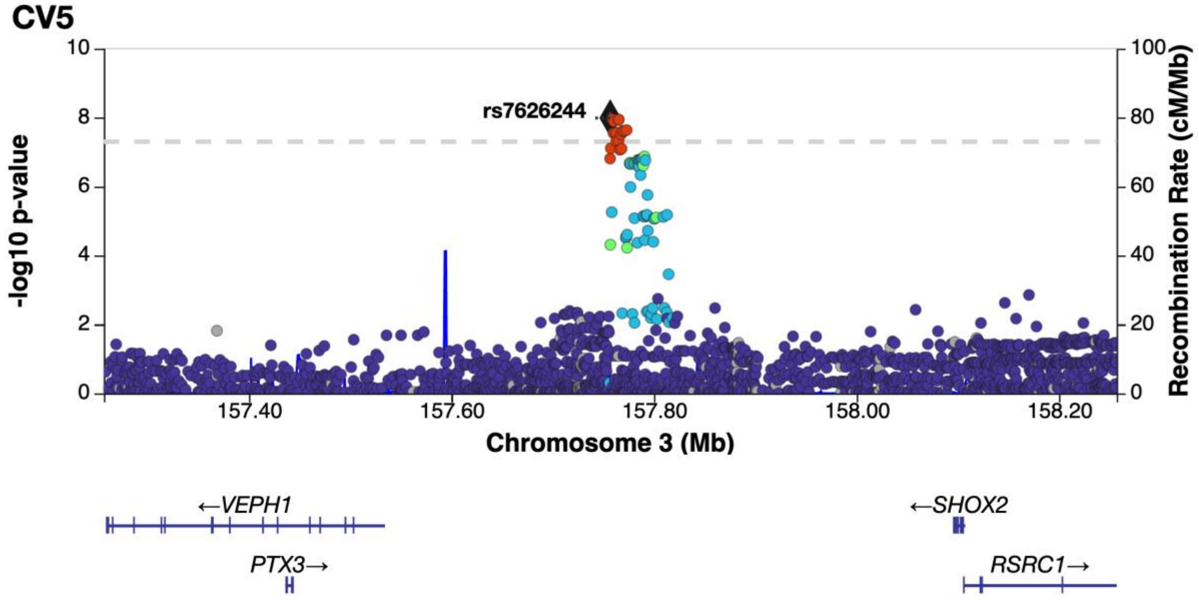

**B**

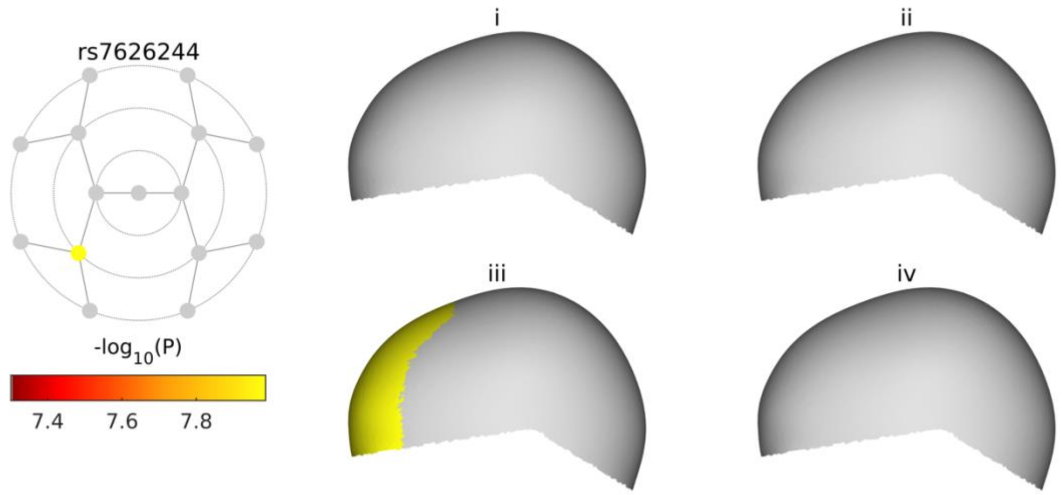

**C**

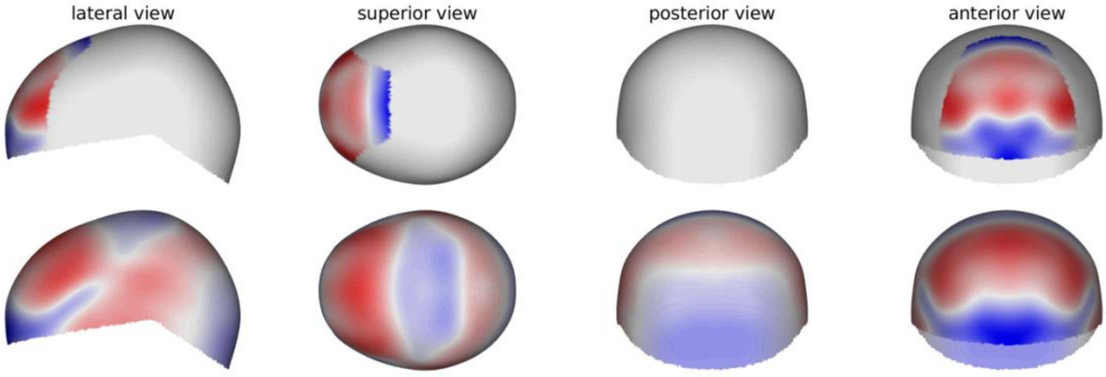

See page 2 for captions

**rs35614773**

**A**

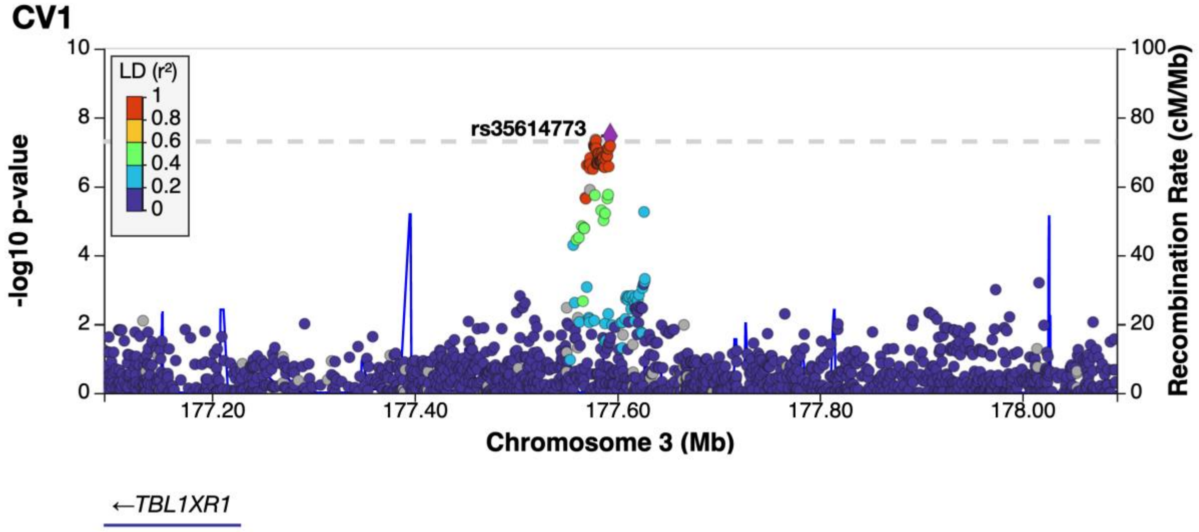

**B**

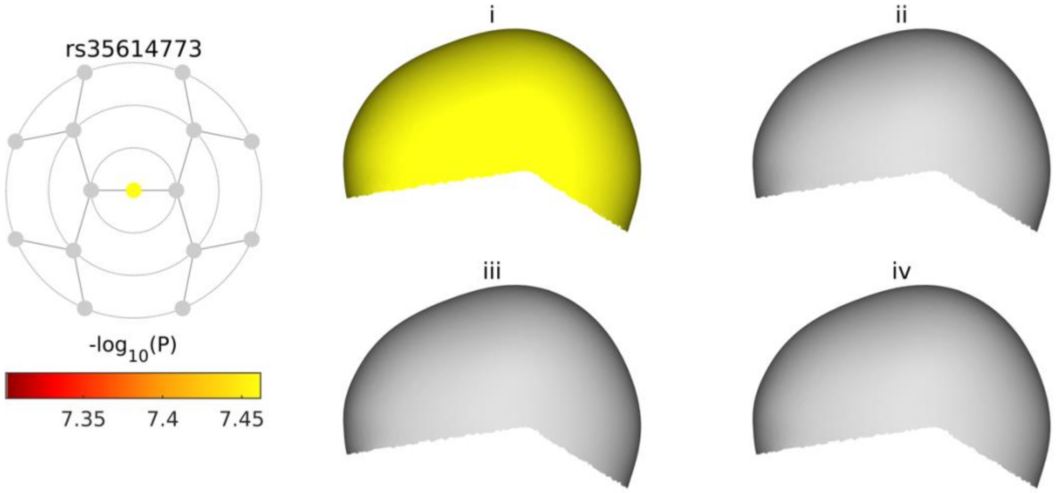

**C**

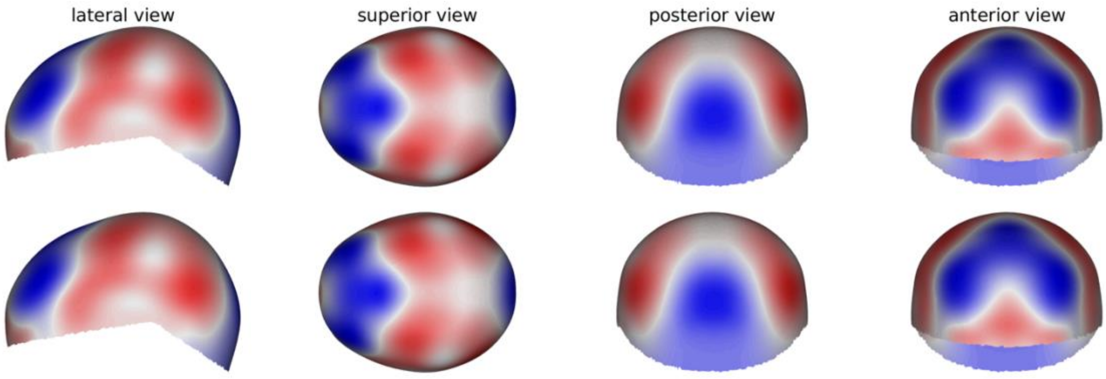

See page 2 for captions

**rs1351637**

**A**

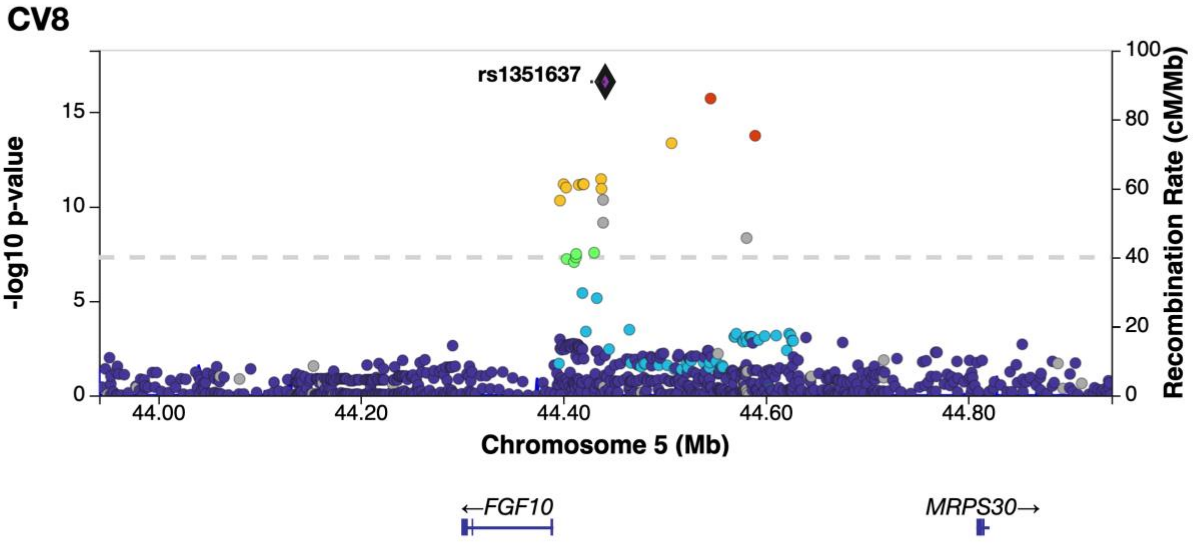

**B**

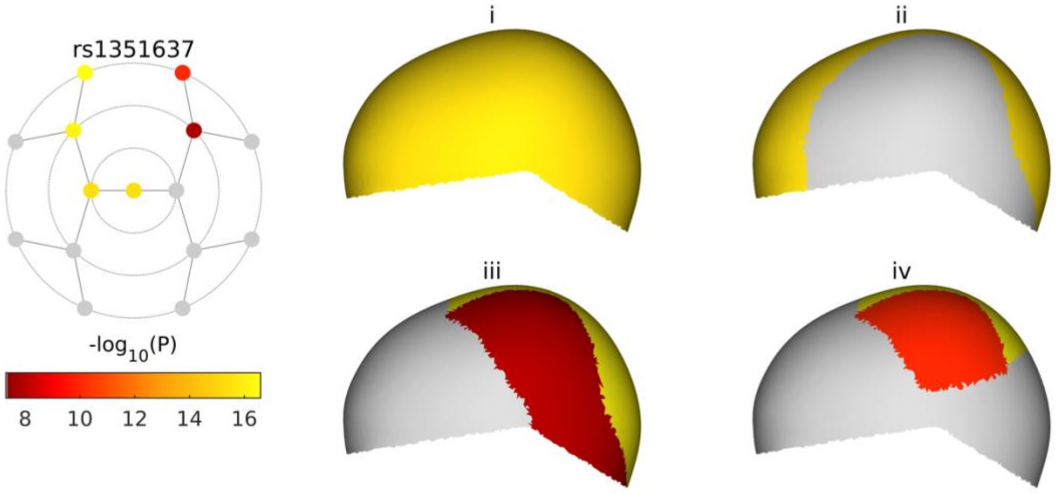

**C**

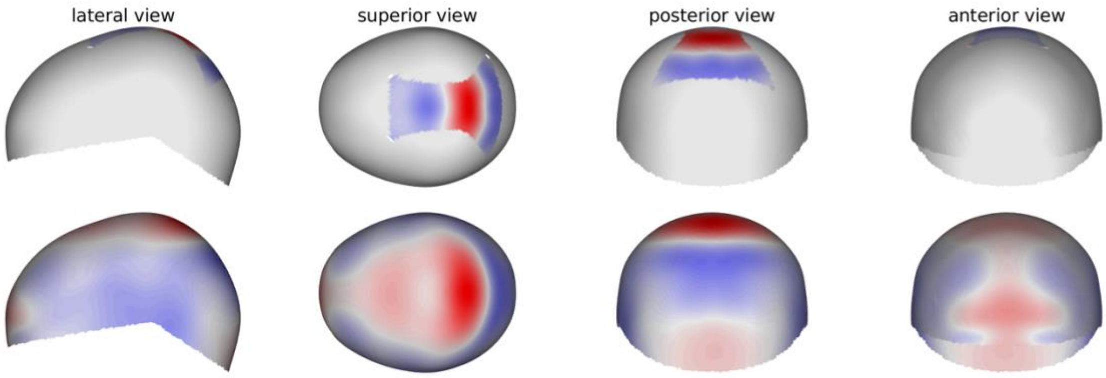

See page 2 for captions

**rs3822730**

**A**

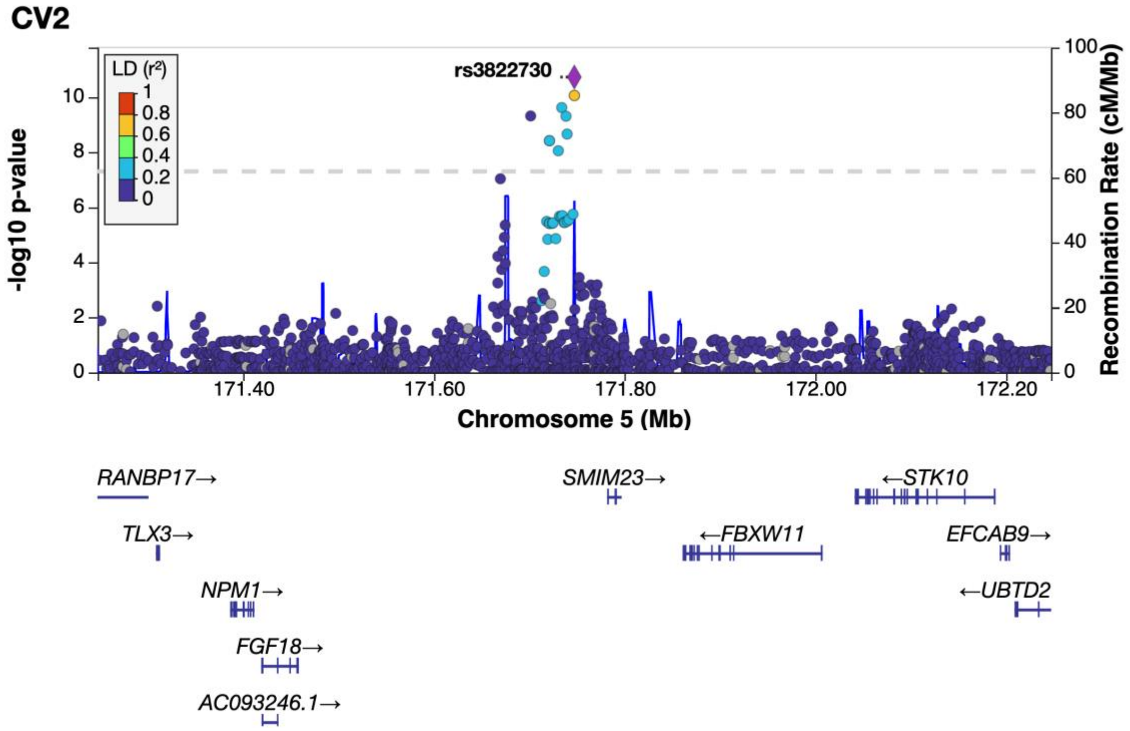

**B**

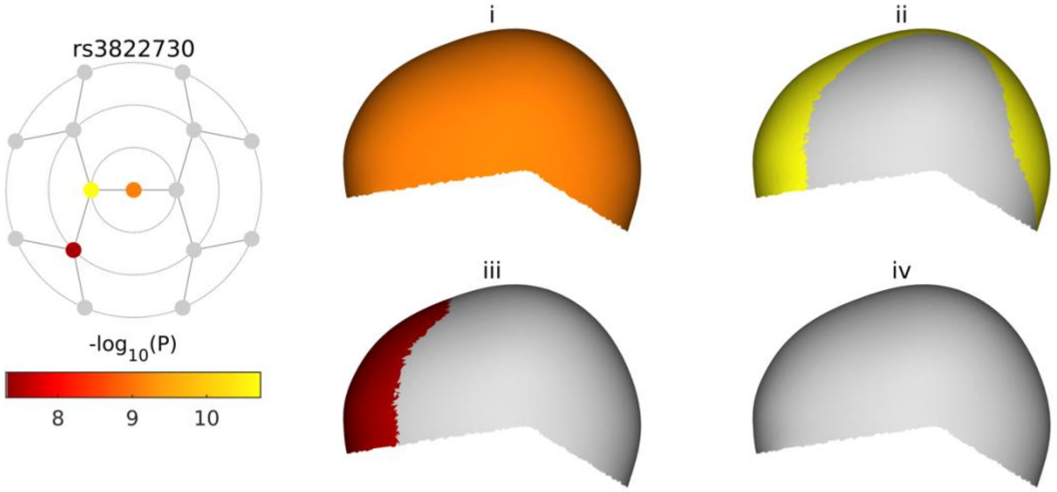

**C**

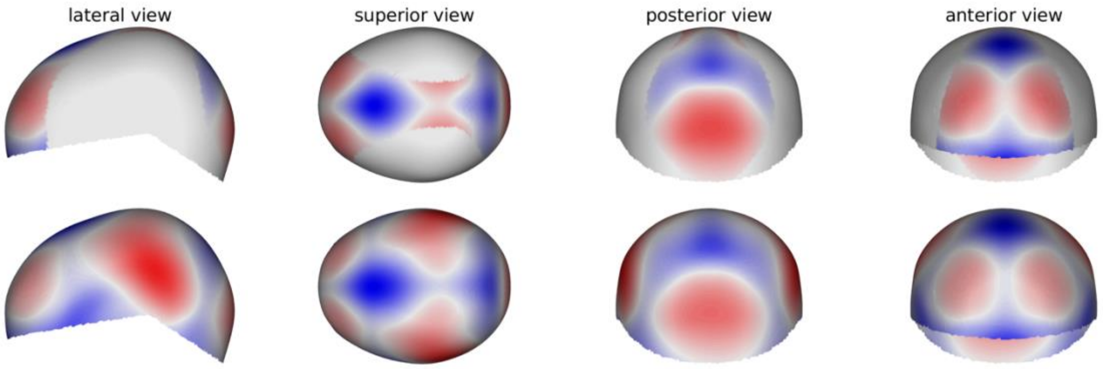

See page 2 for captions

**rs4714260**

**A**

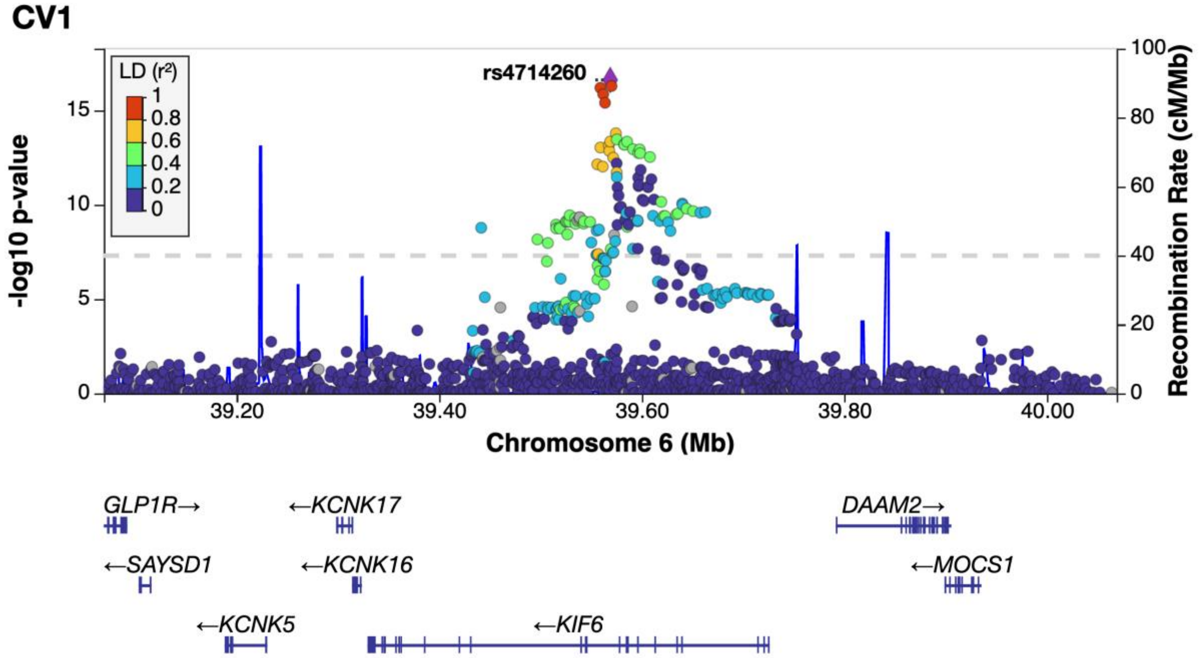

**B**

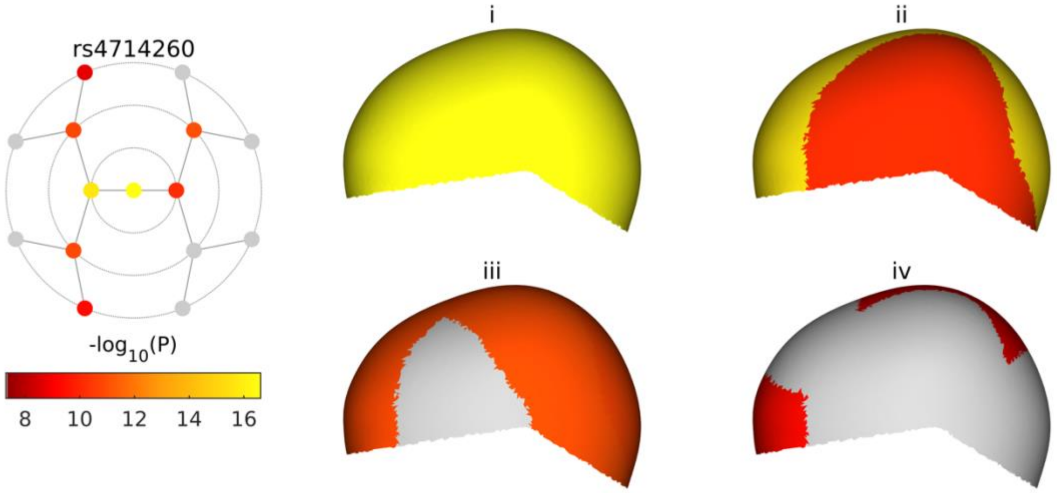

**C**

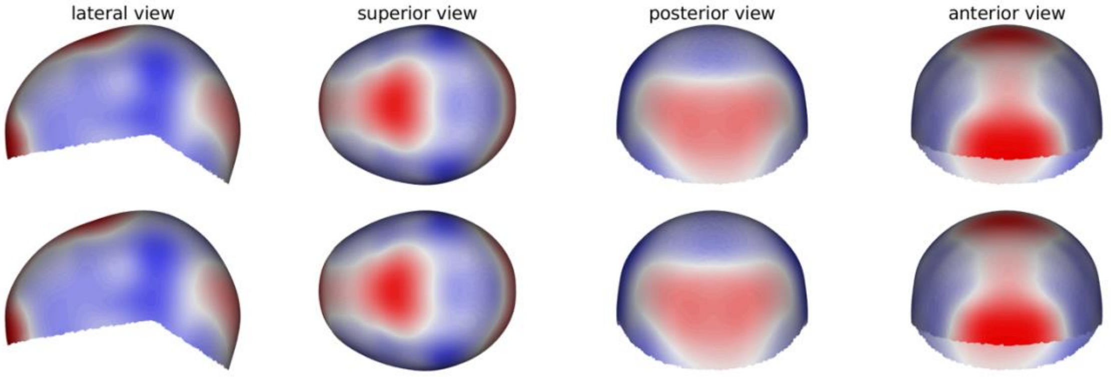

See page 2 for captions

**rs3799970**

**A**

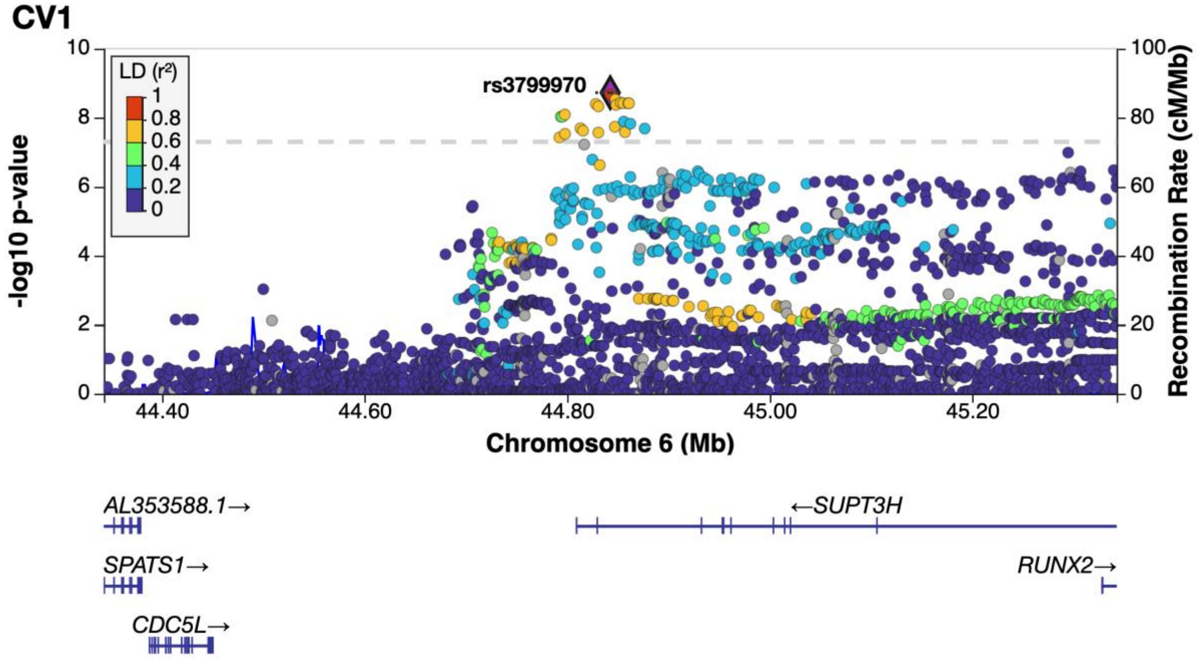

**B**

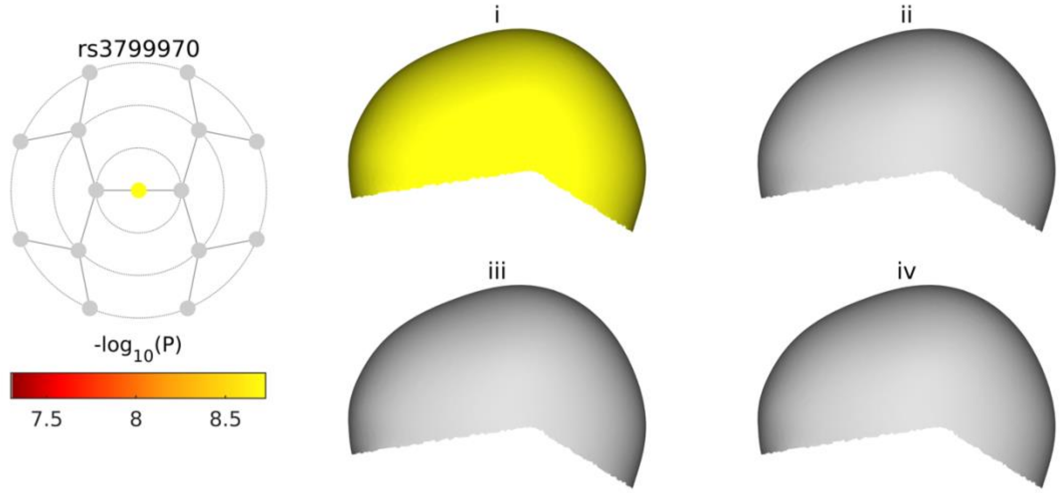

**C**

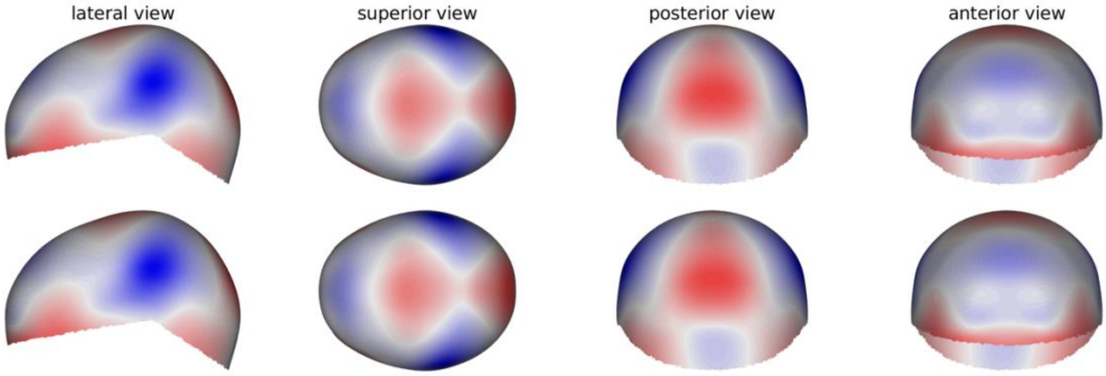

See page 2 for captions

**rs9491697**

**A**

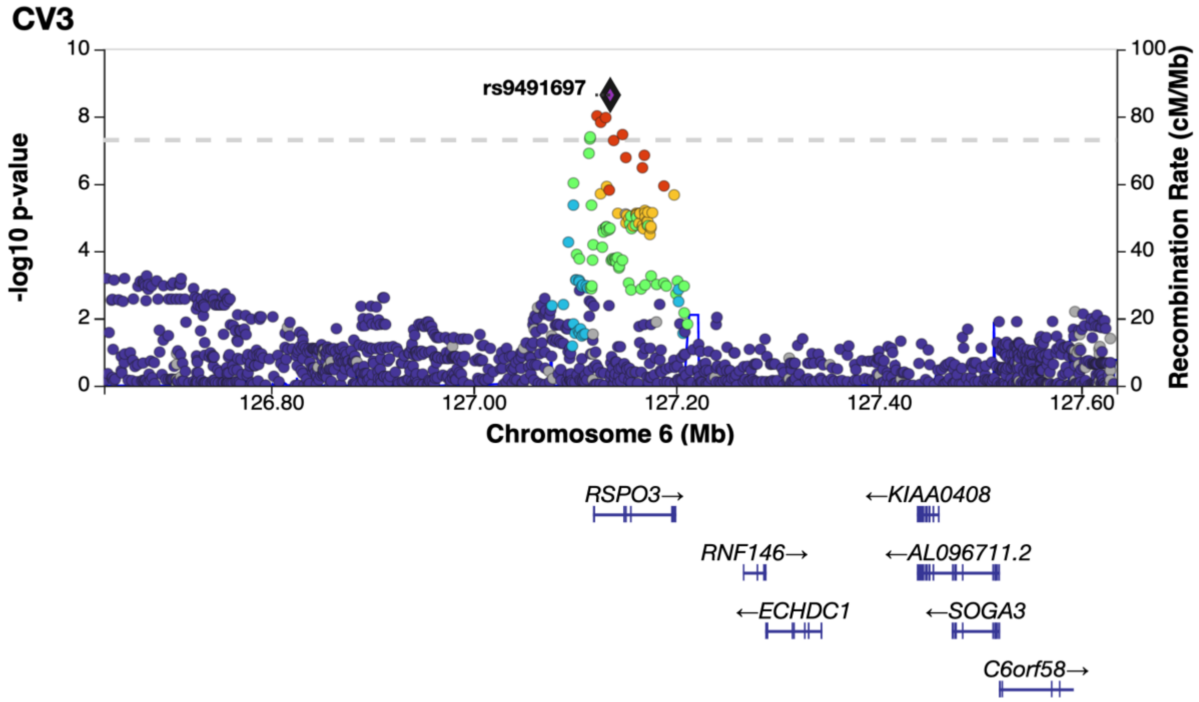

**B**

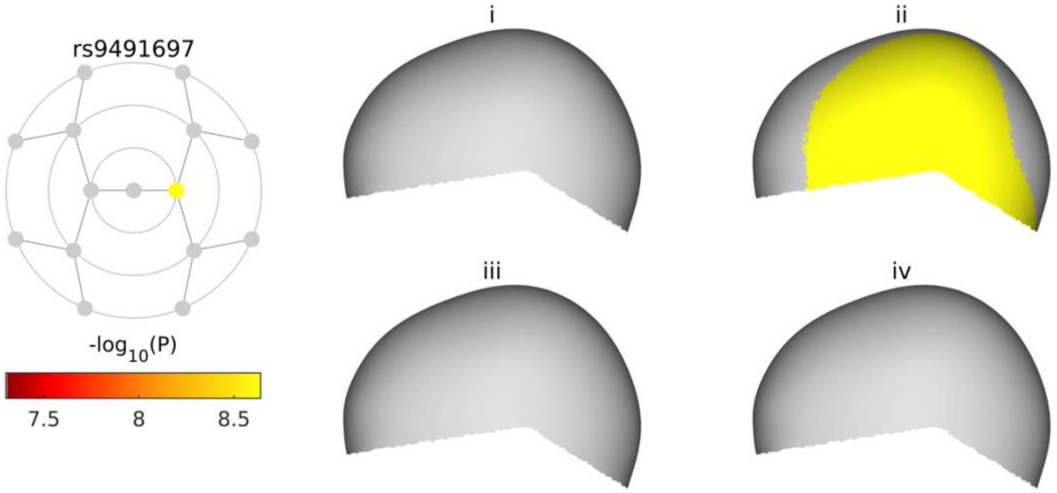

**C**

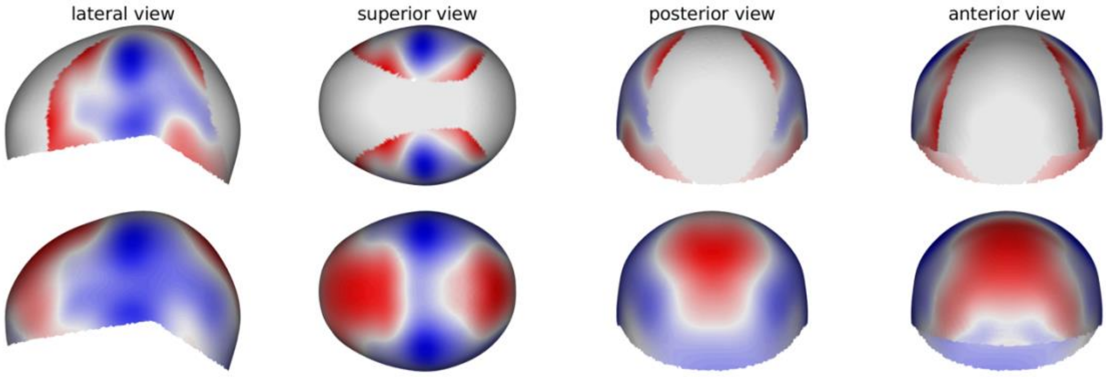

See page 2 for captions

**rs296418**

**A**

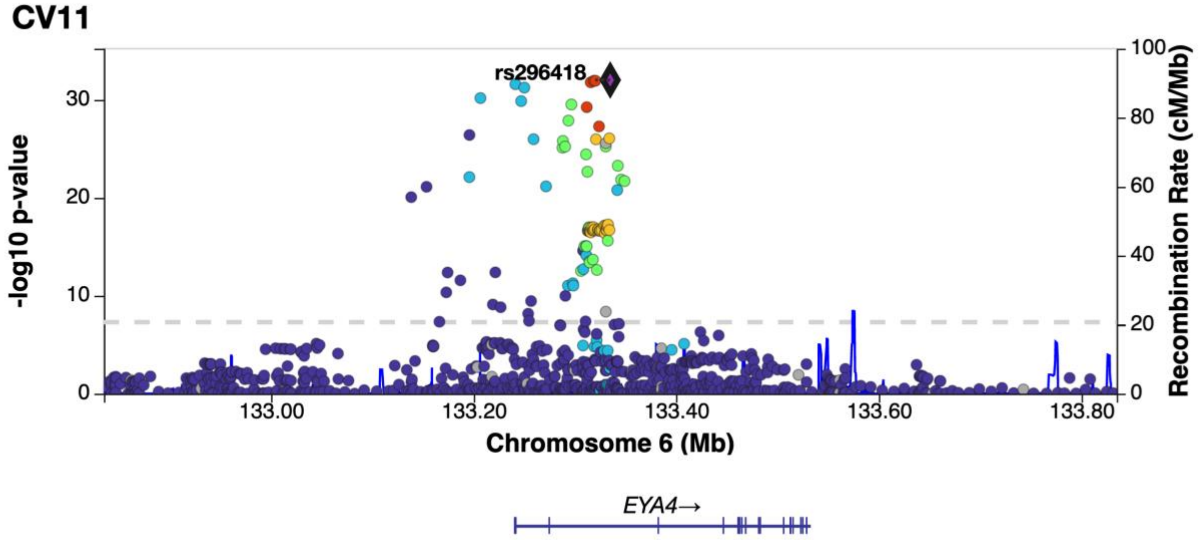

**B**

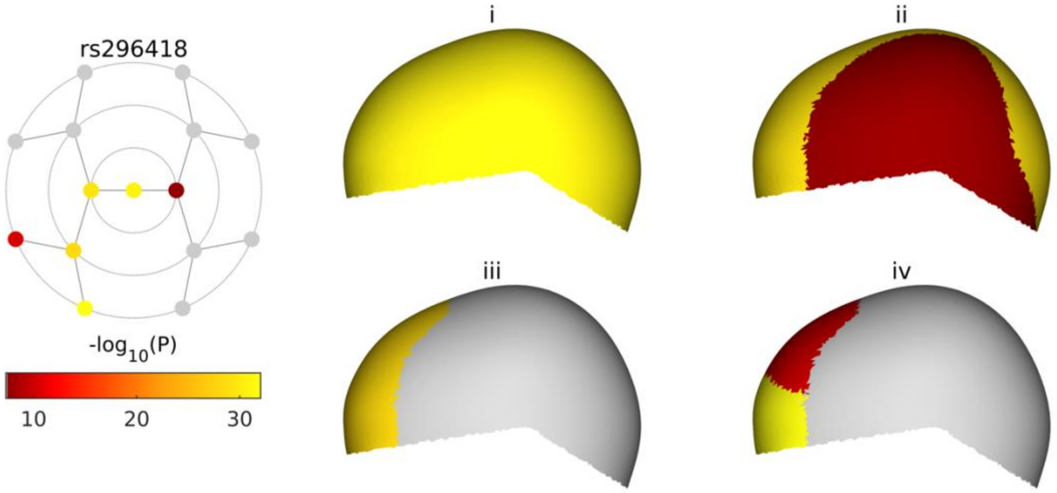

**C**

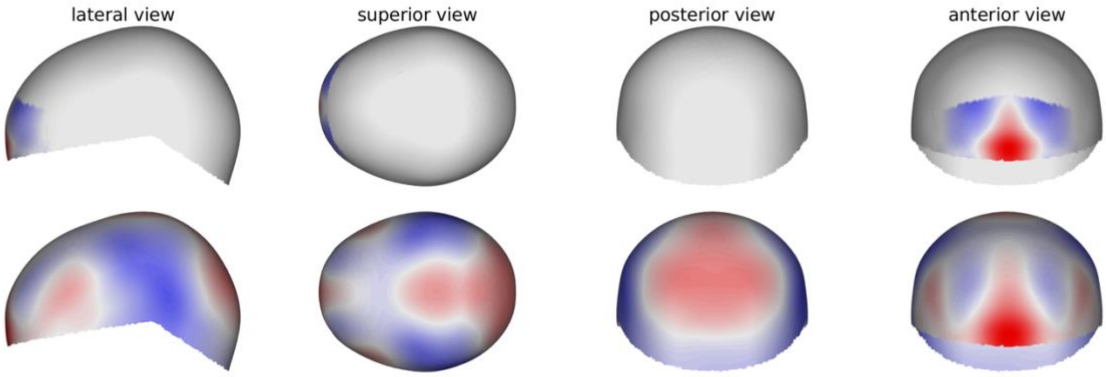

See page 2 for captions

**rs148673350**

**A**

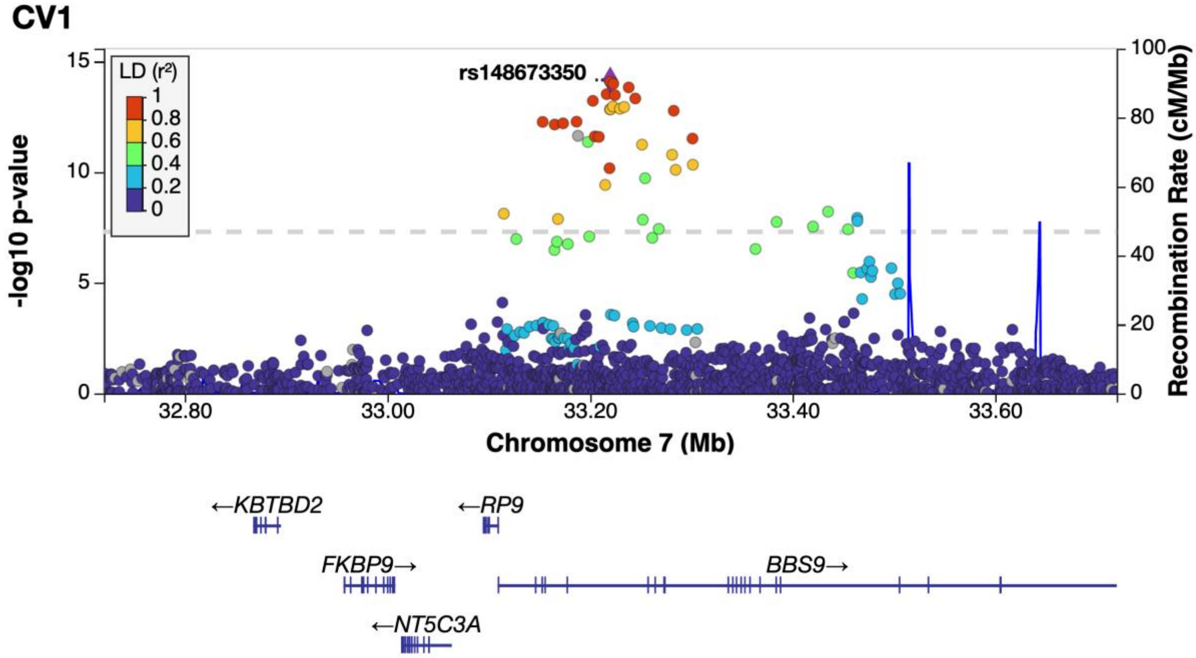

**B**

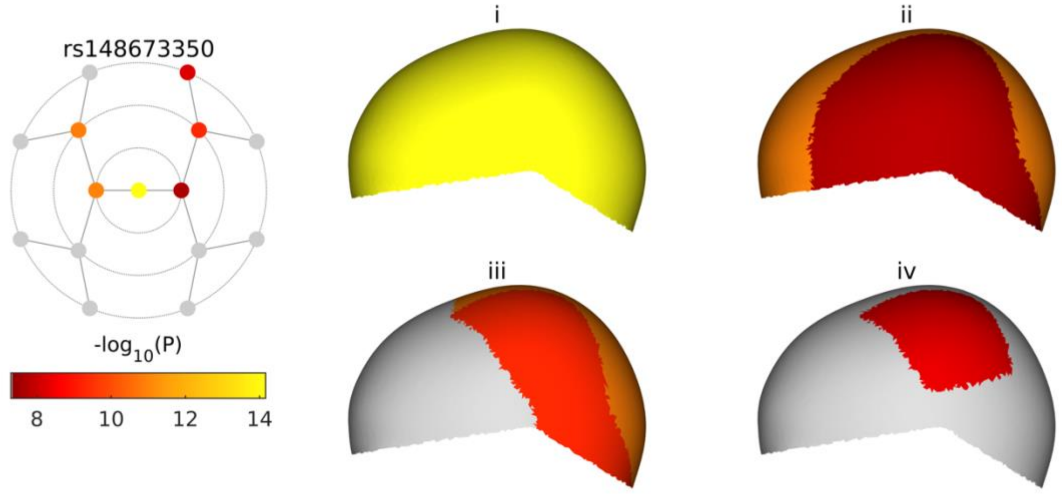

**C**

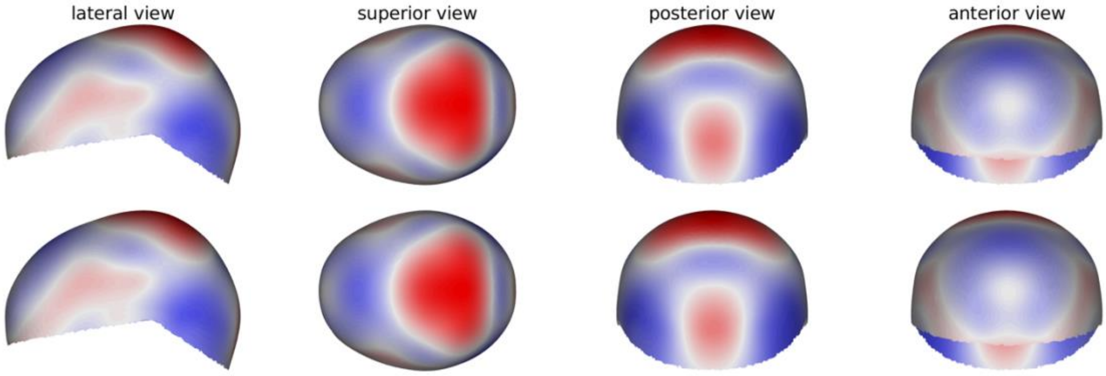

See page 2 for captions

**rs202055590**

**A**

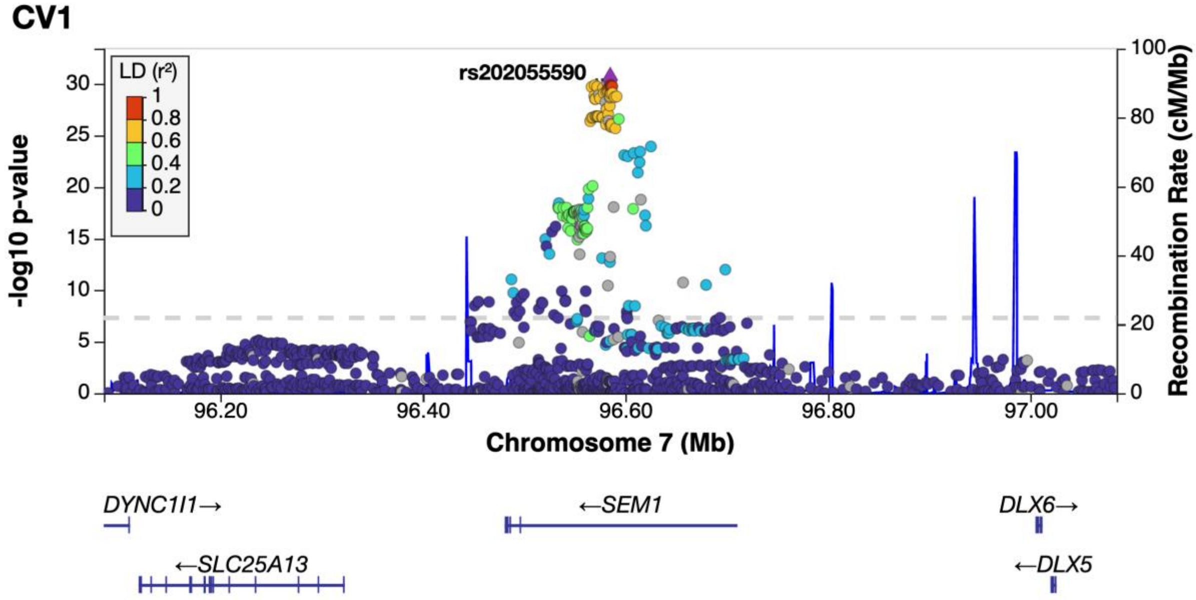

**B**

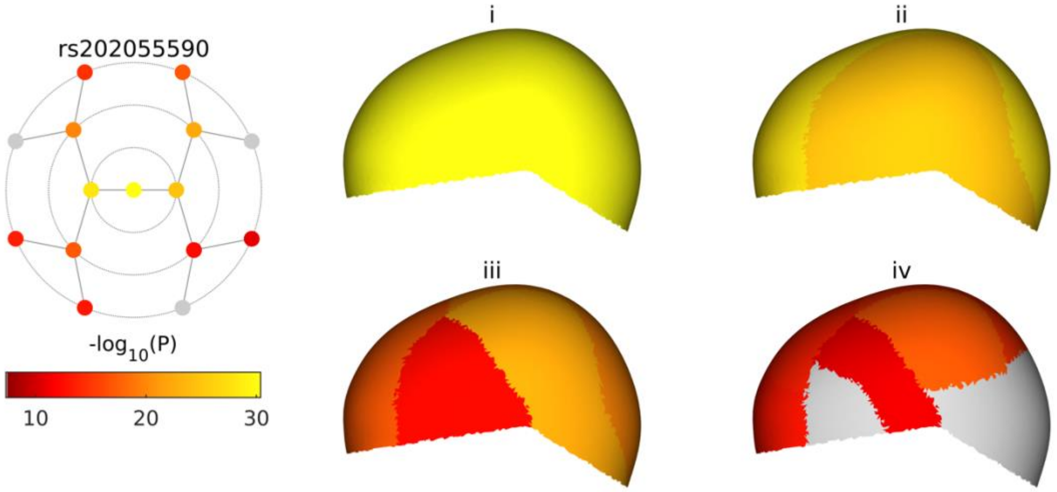

**C**

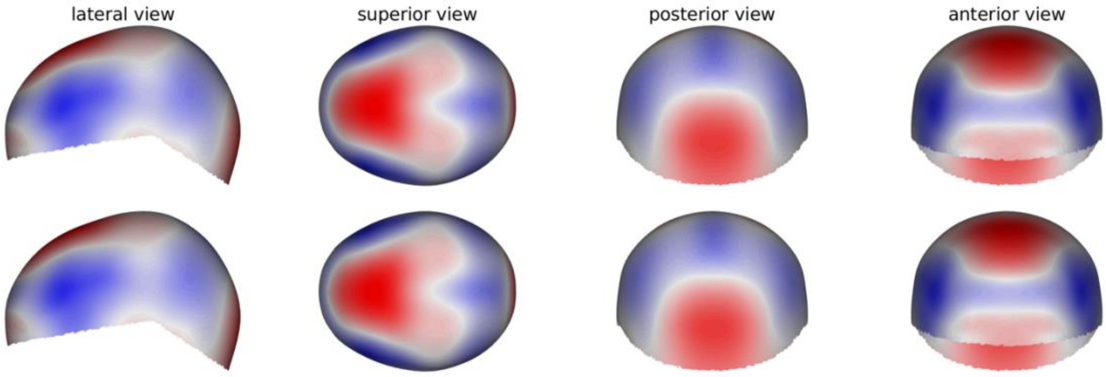

See page 2 for captions

**rs1581525**

**A**

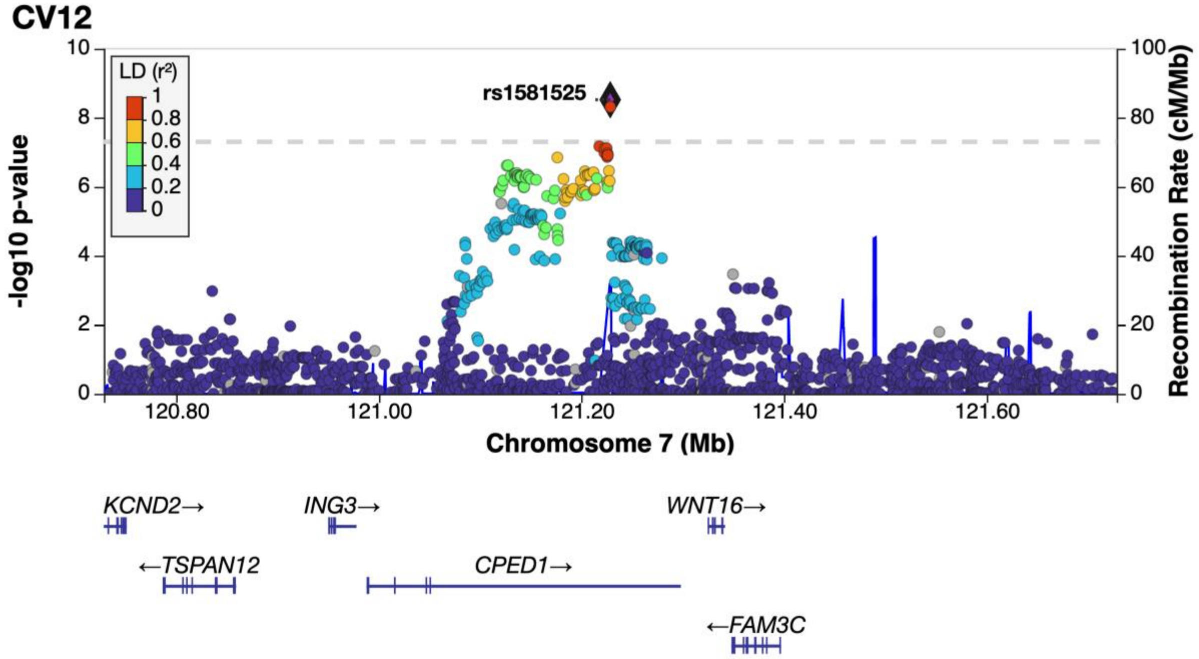

**B**

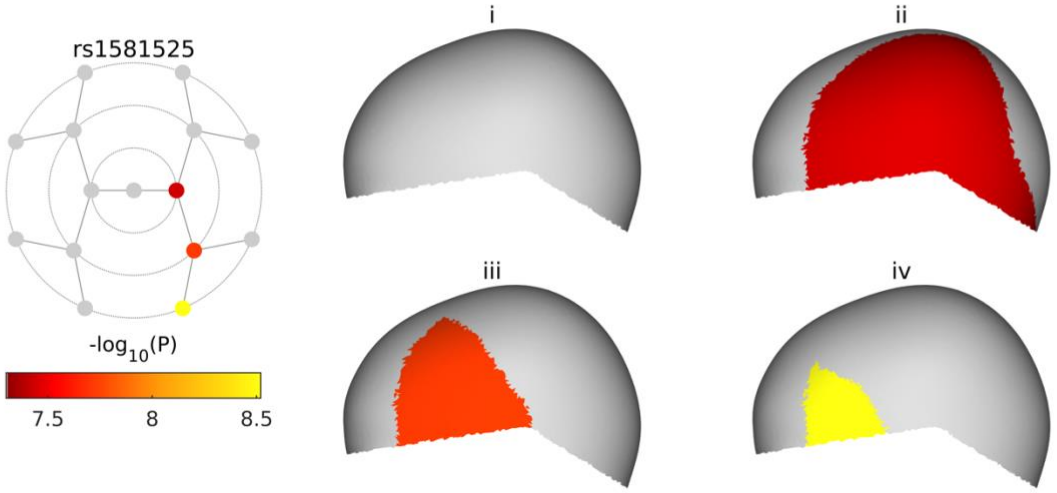

**C**

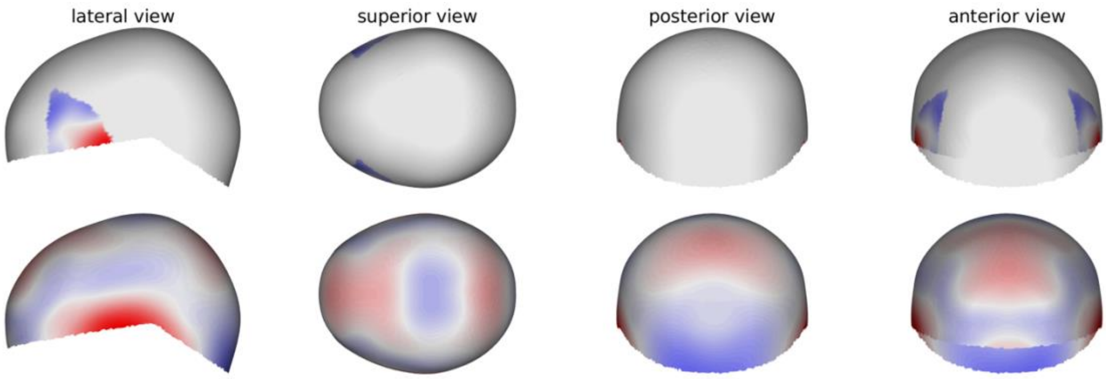

See page 2 for captions

**rs147676525**

**A**

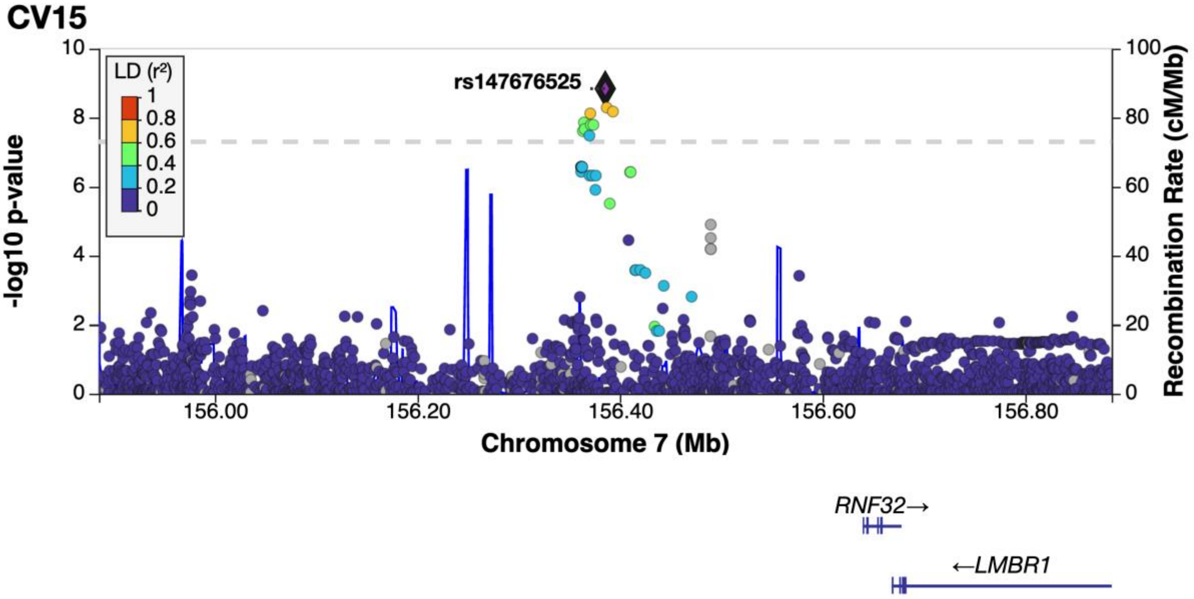

**B**

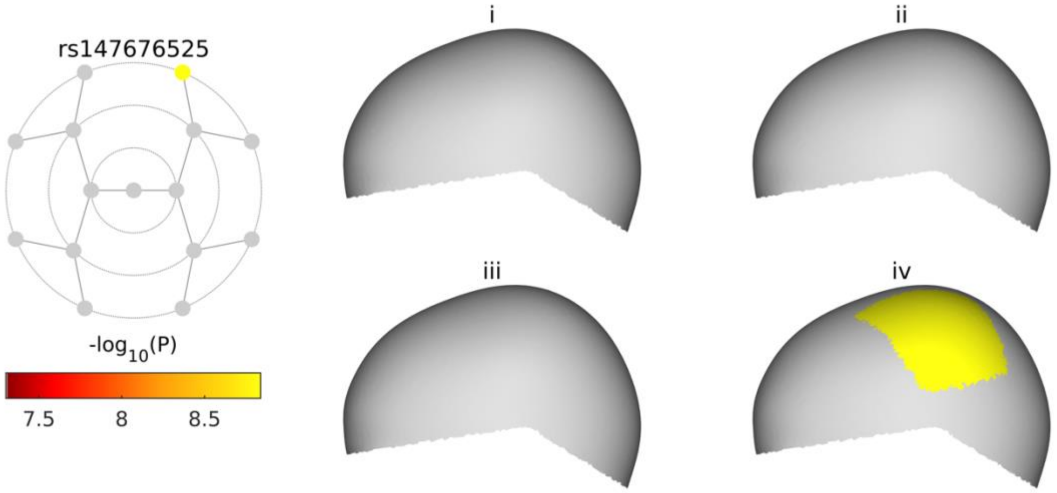

**C**

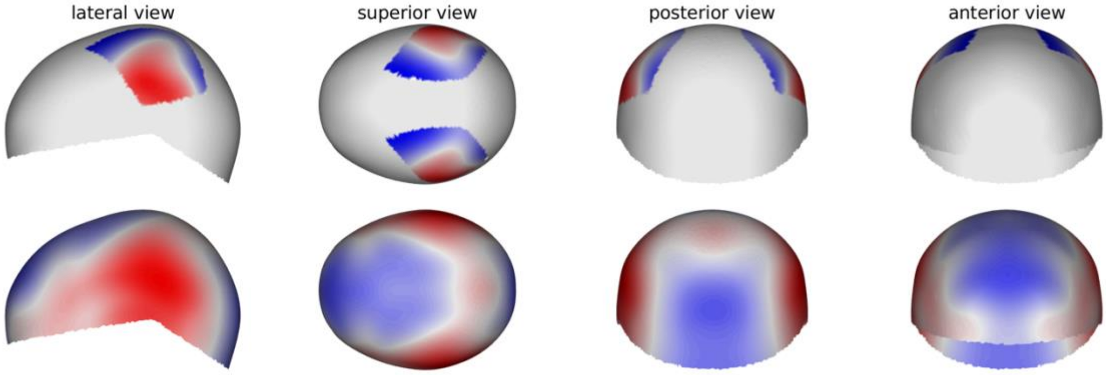

See page 2 for captions

**rs7813717**

**A**

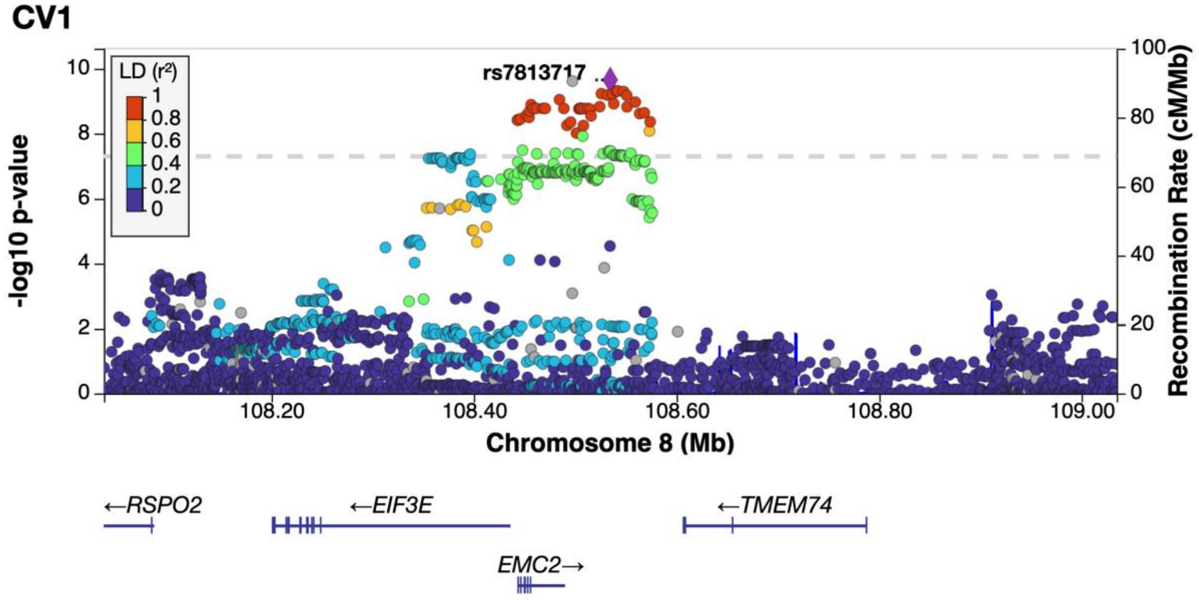

**B**

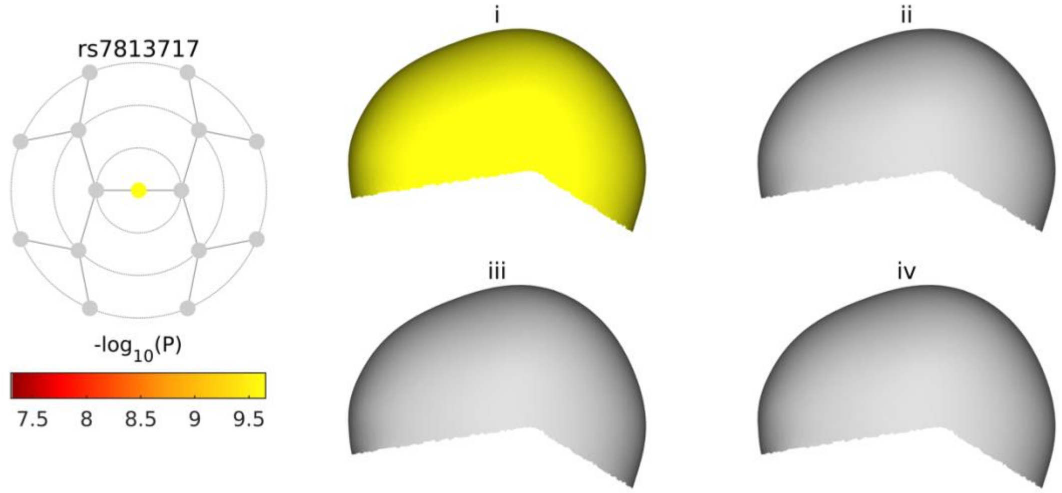

**C**

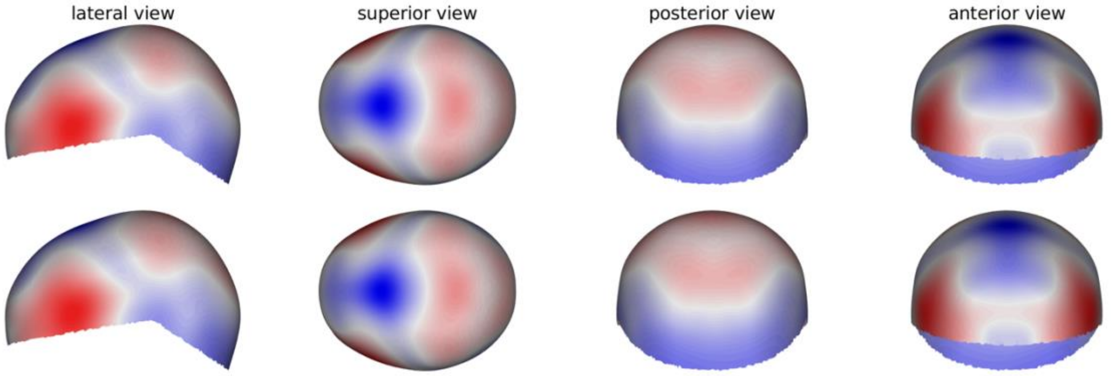

See page 2 for captions

**rs10120728**

**A**

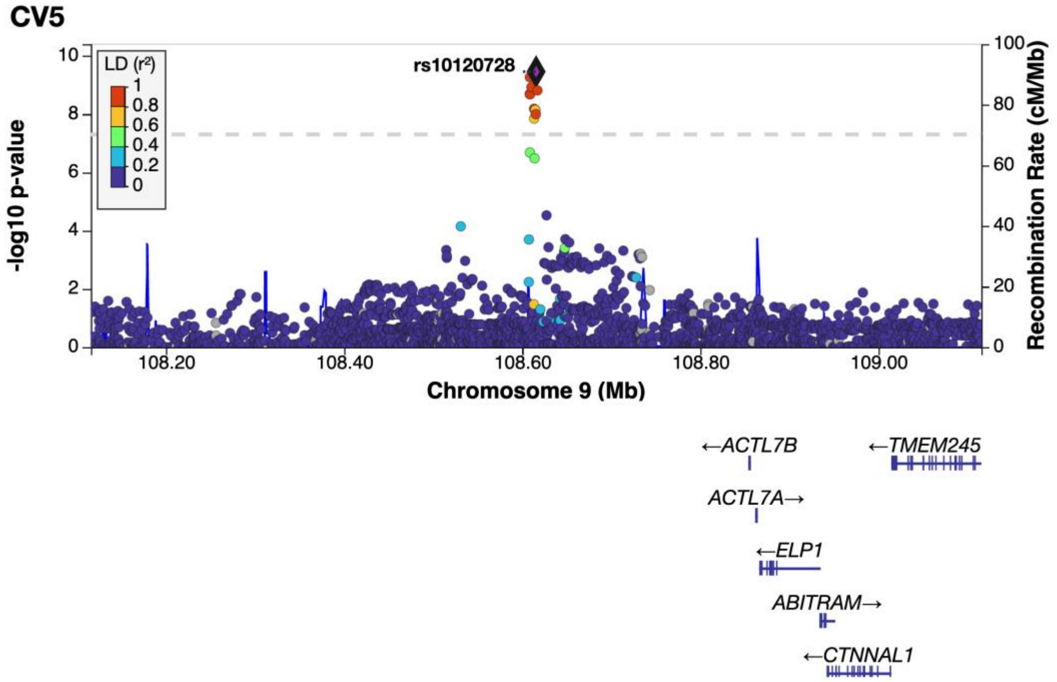

**B**

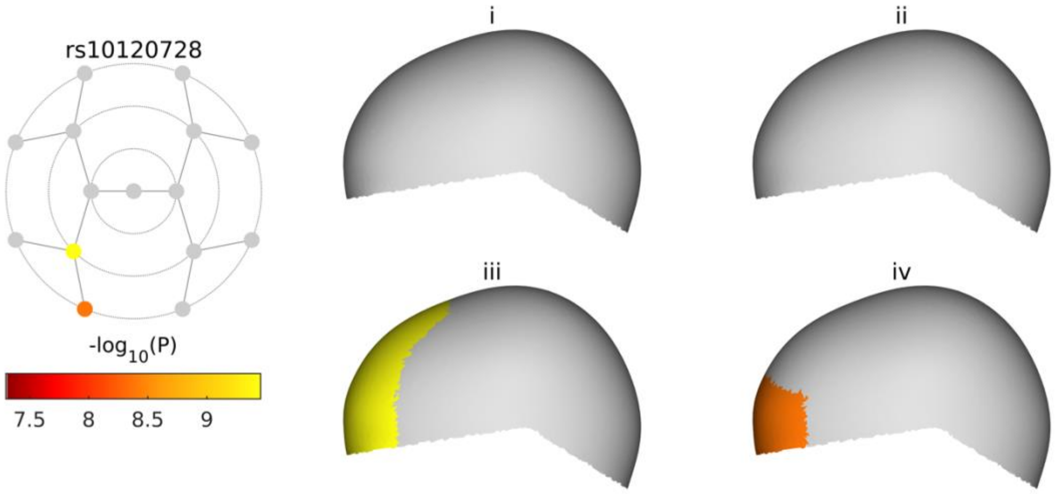

**C**

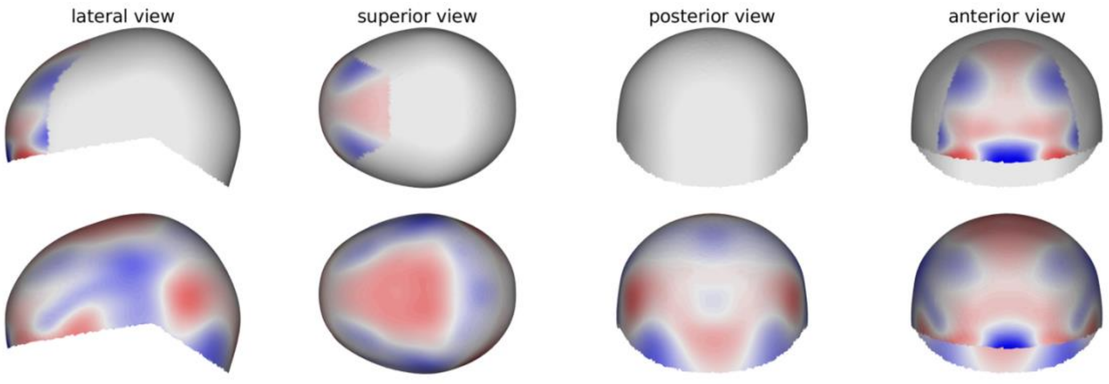

See page 2 for captions

**rs7920484**

**A**

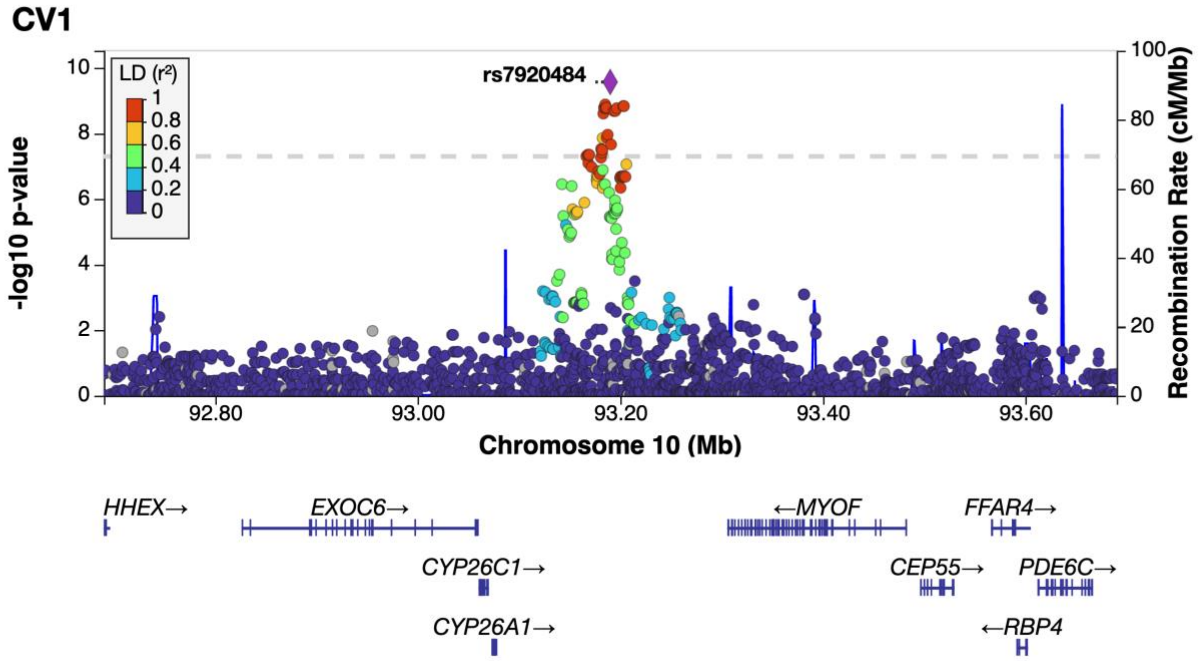

**B**

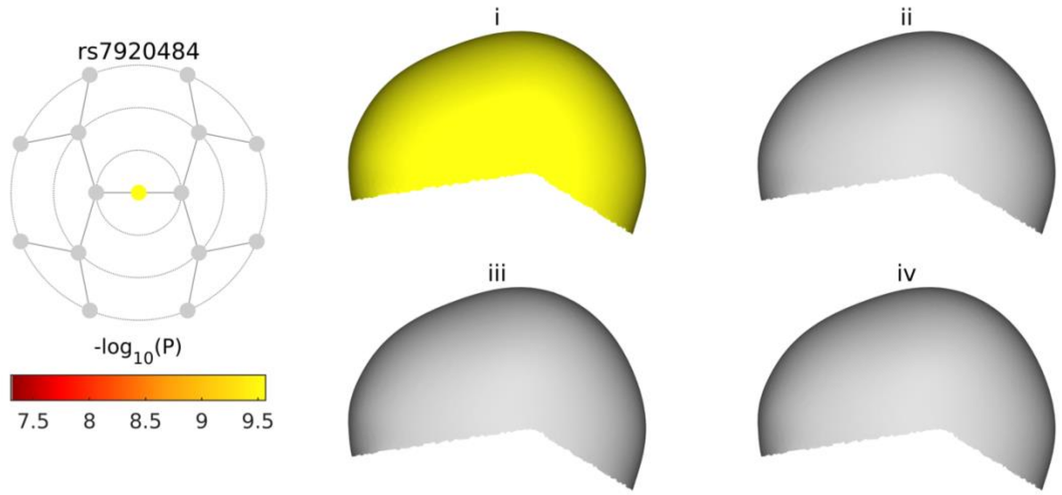

**C**

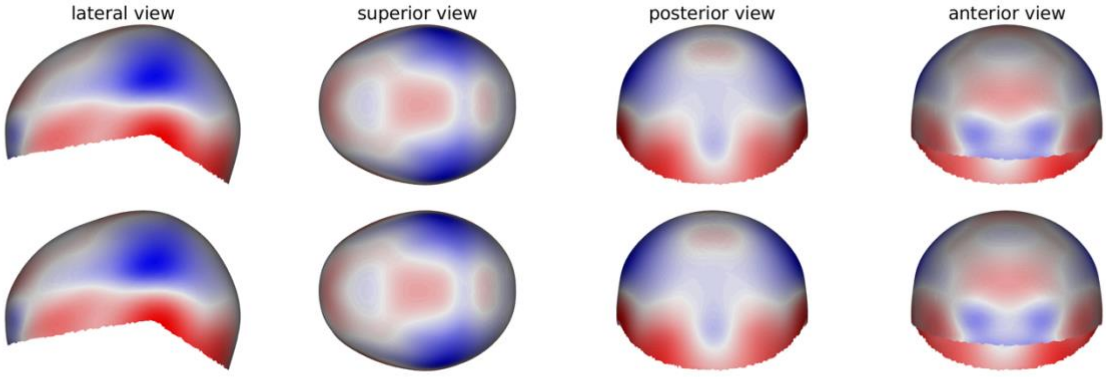

See page 2 for captions

**rs61920200**

**A**

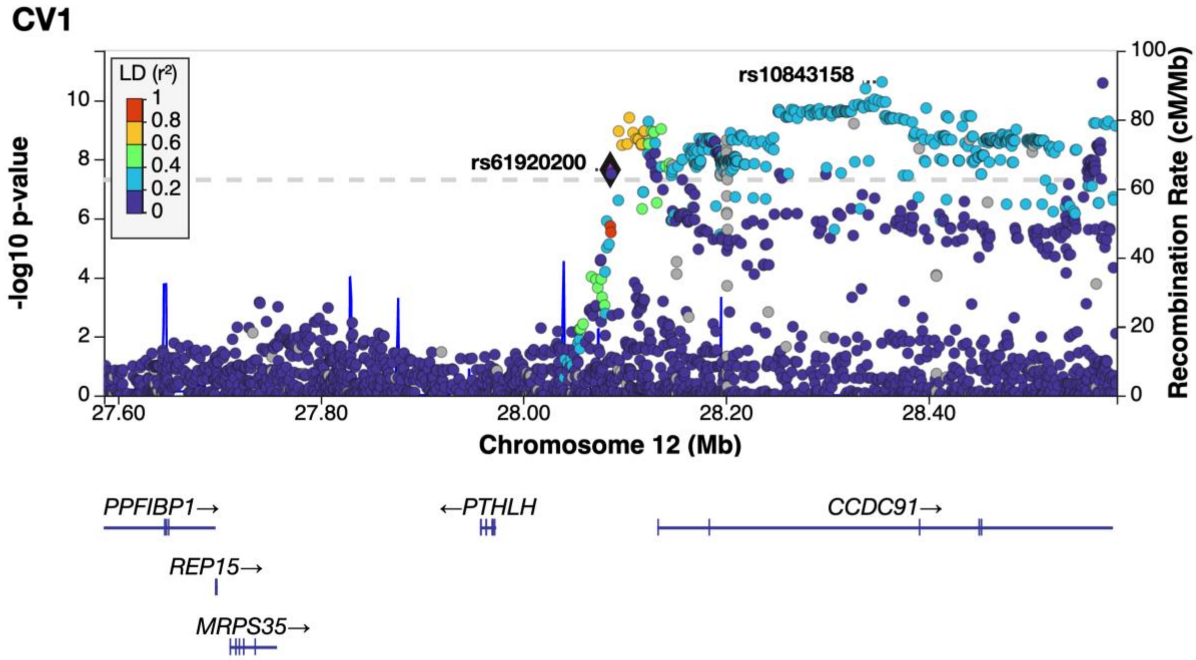

**B**

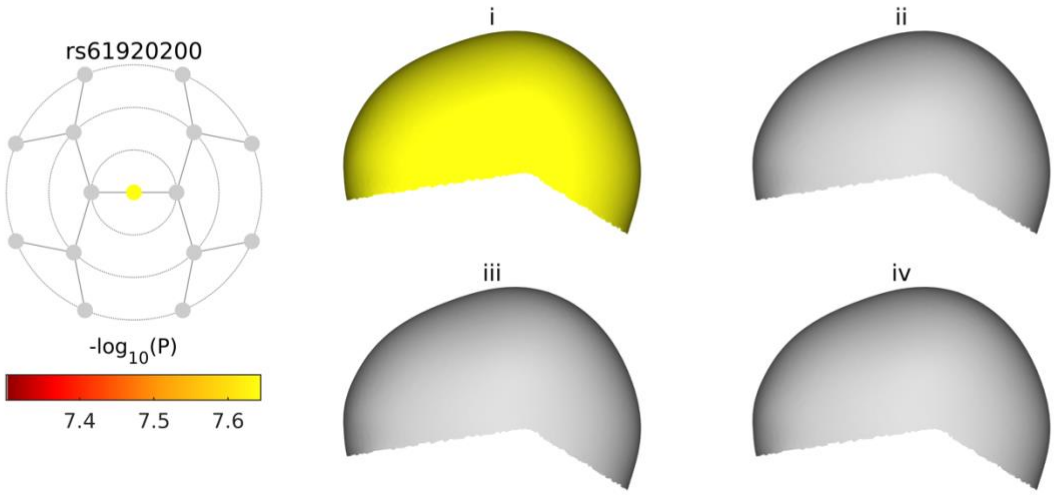

**C**

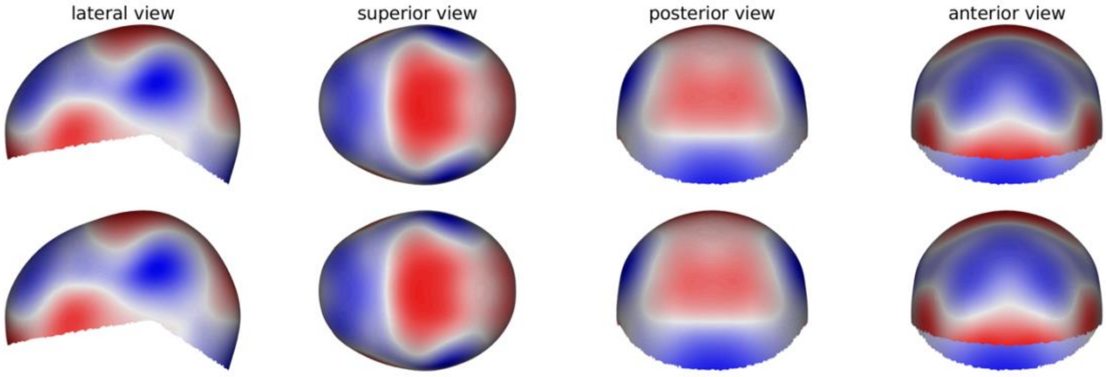

See page 2 for captions

**rs10843158**

**A**

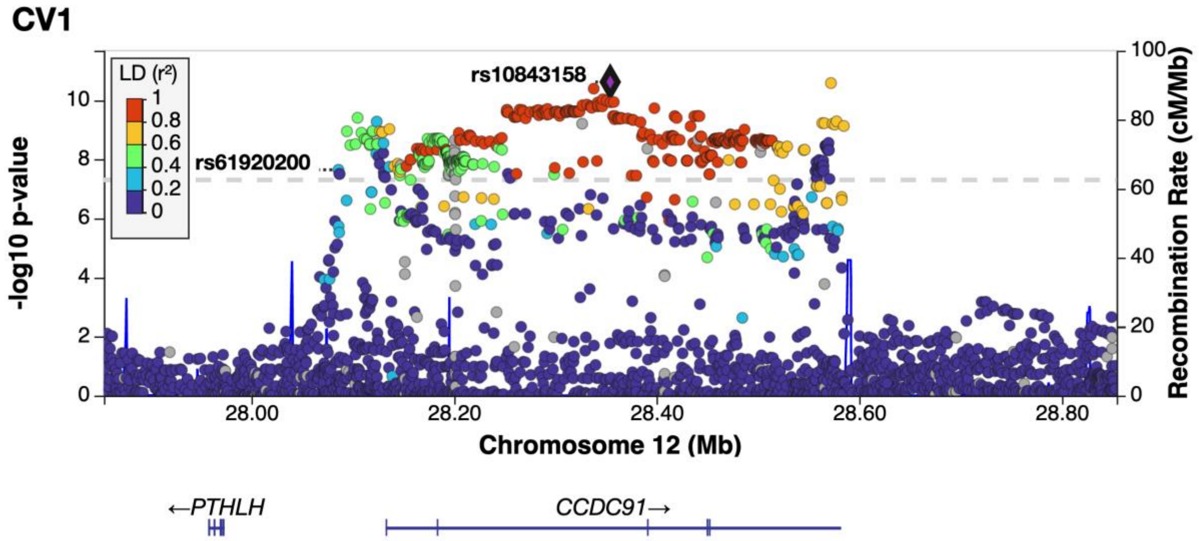

**B**

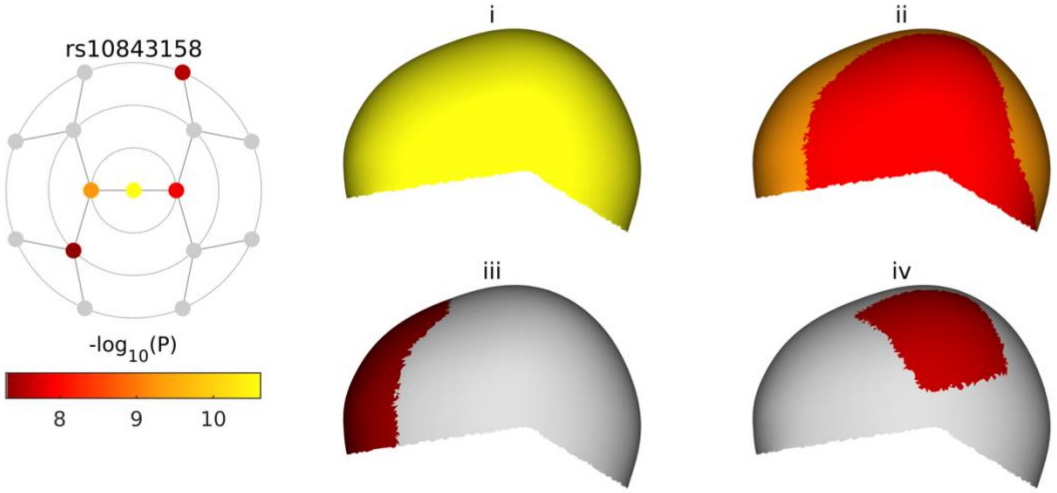

**C**

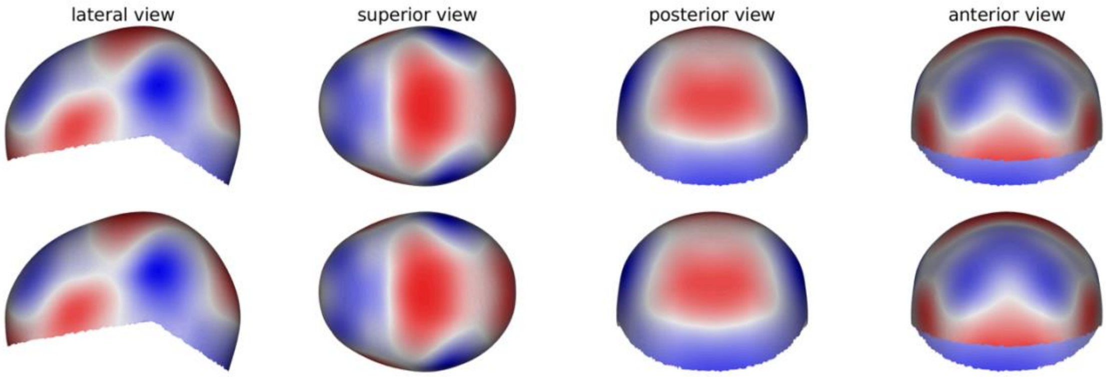

See page 2 for captions

**rs151174669**

**A**

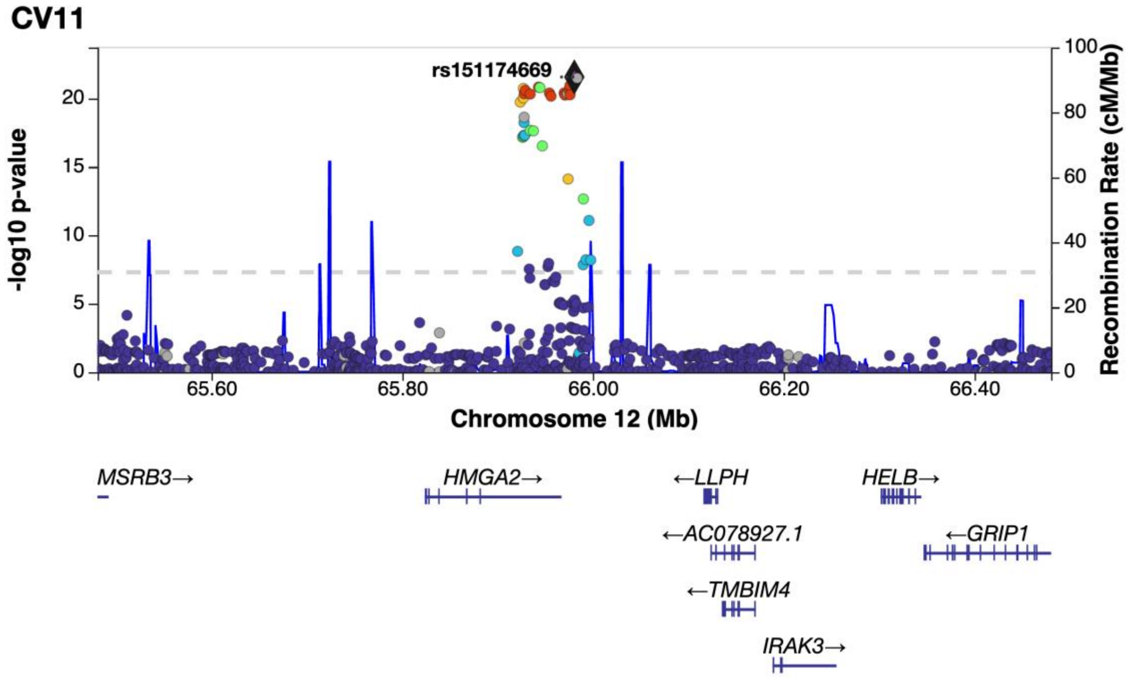

**B**

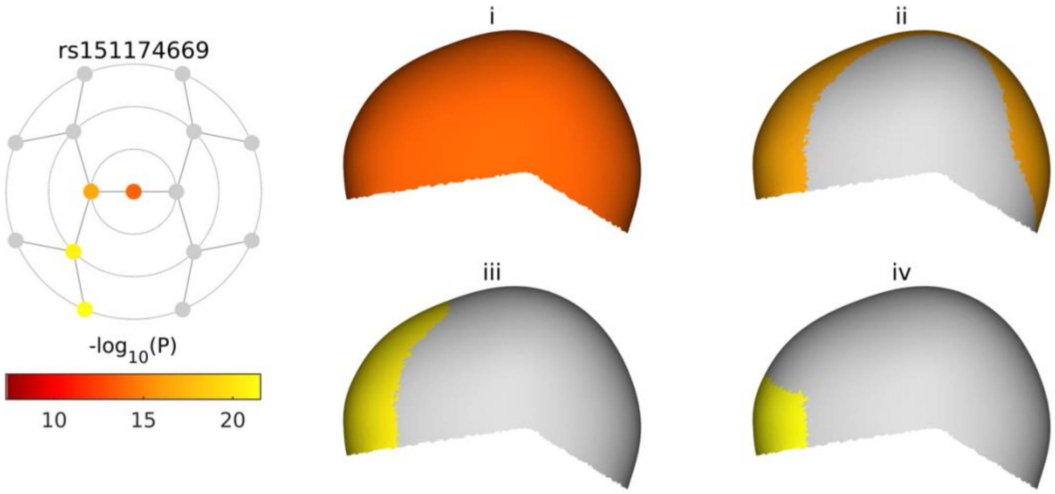

**C**

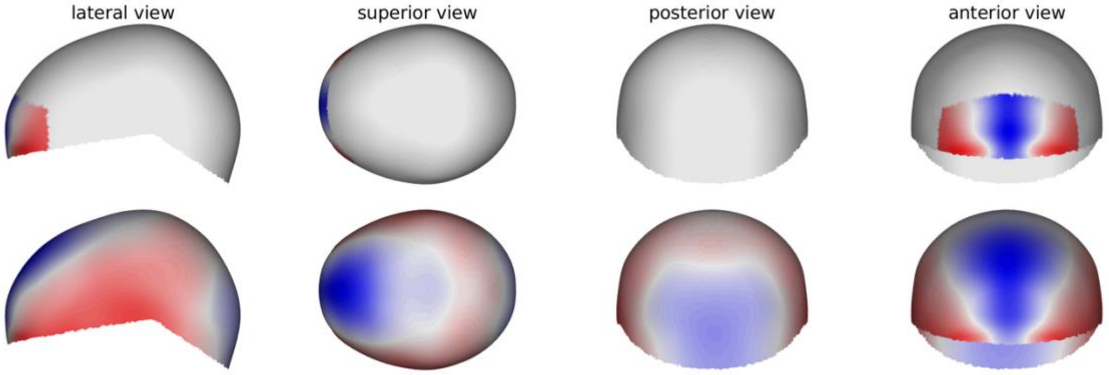

See page 2 for captions

**rs11609649**

**A**

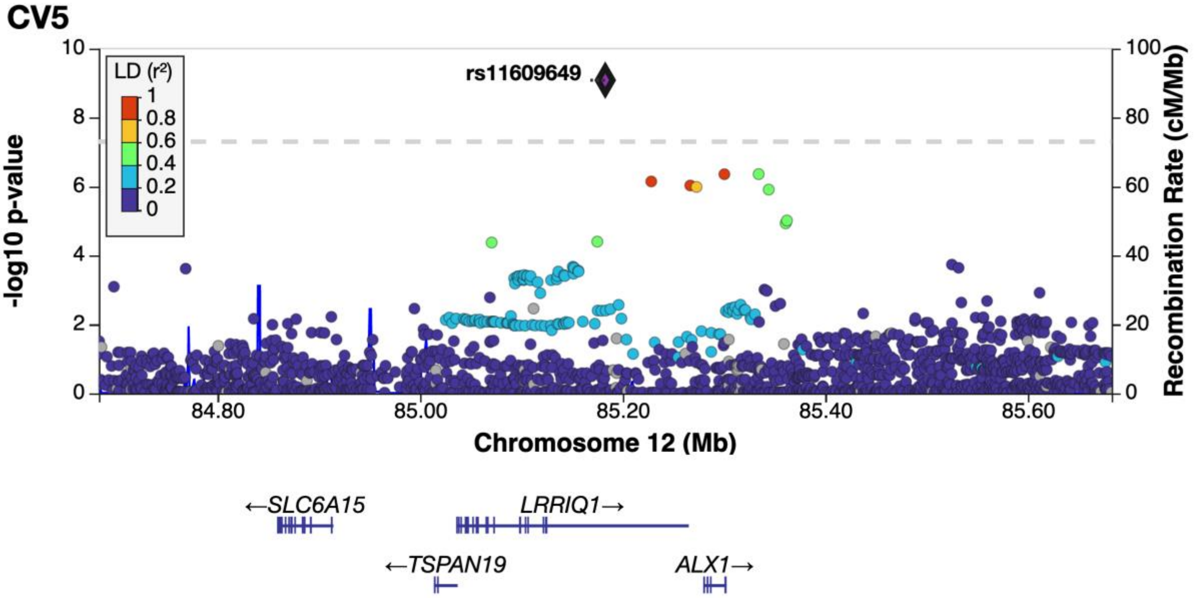

**B**

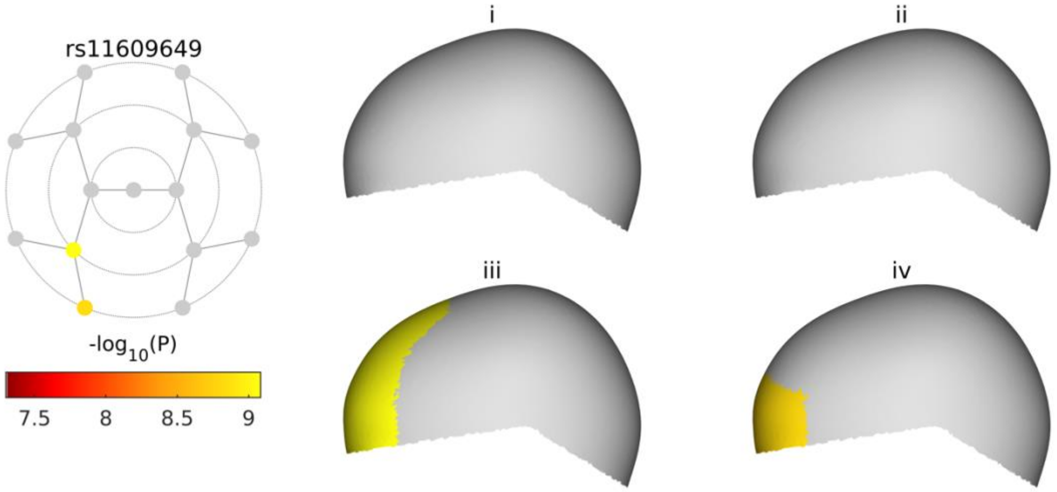

**C**

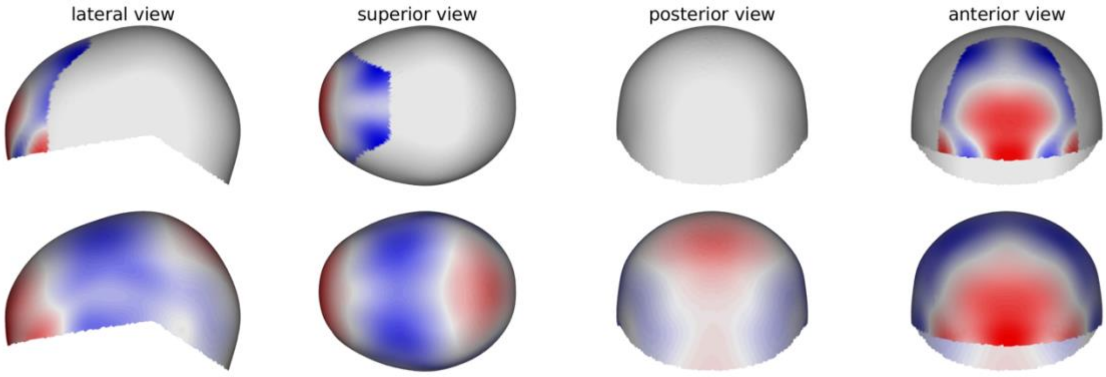

See page 2 for captions

**rs1034266**

**A**

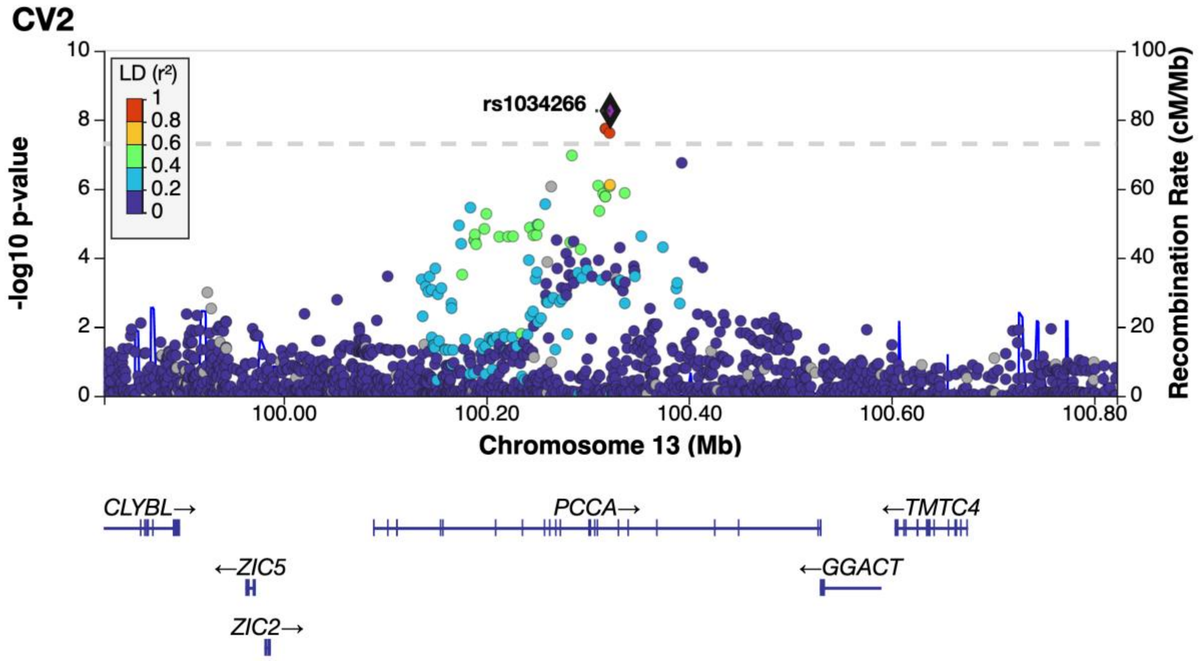

**B**

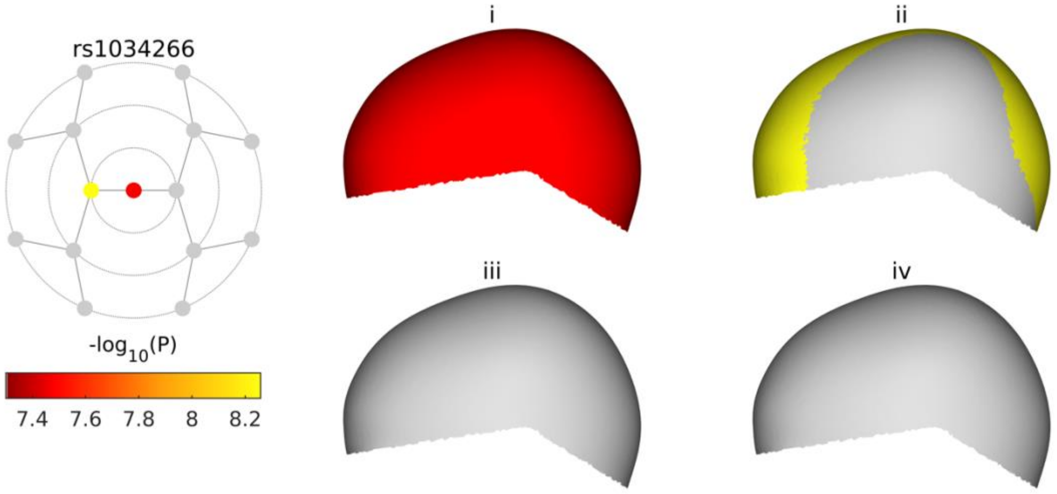

**C**

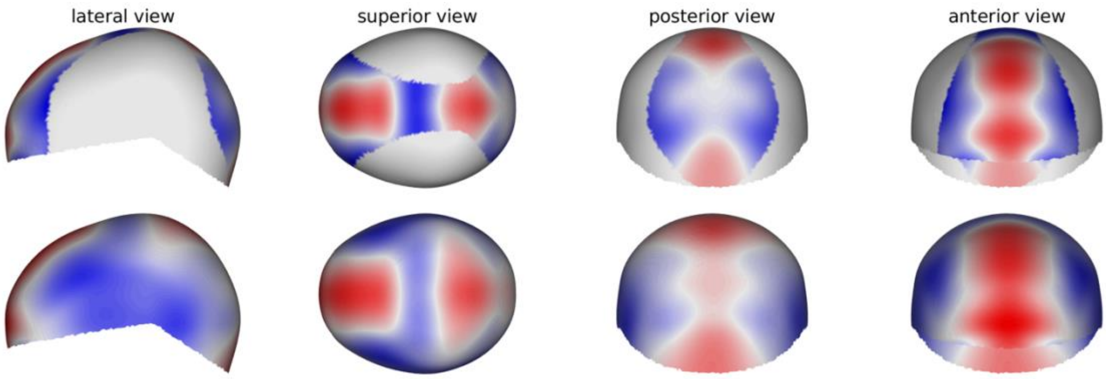

See page 2 for captions

**rs1380208**

**A**

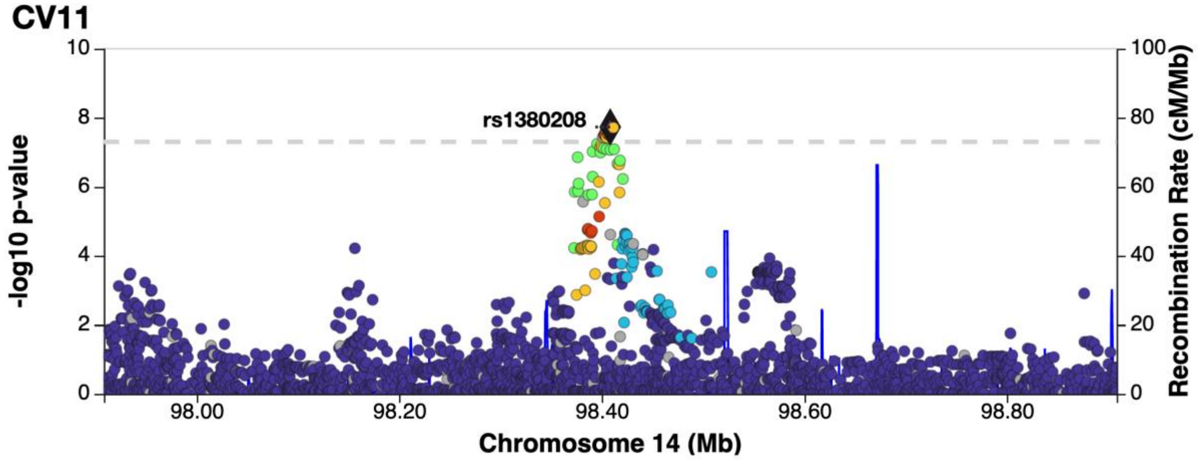

**B**

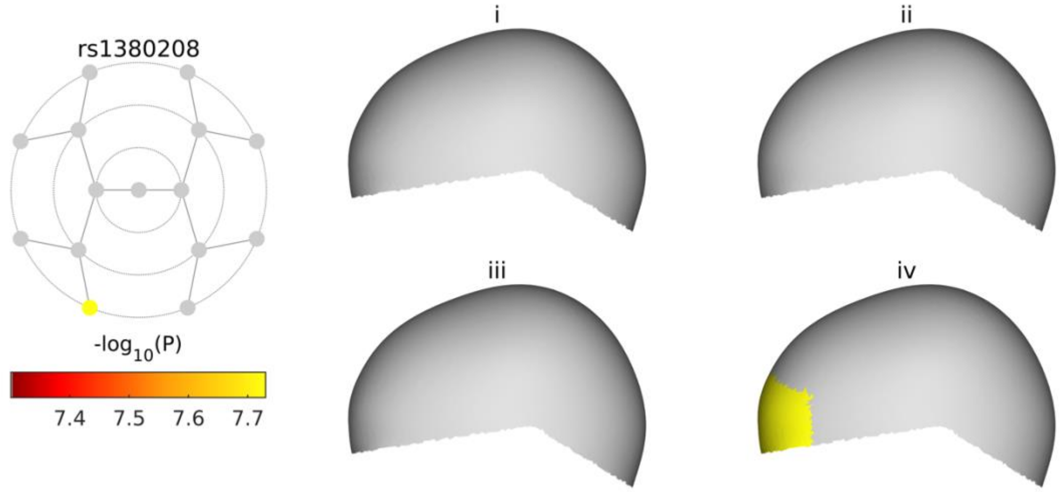

**C**

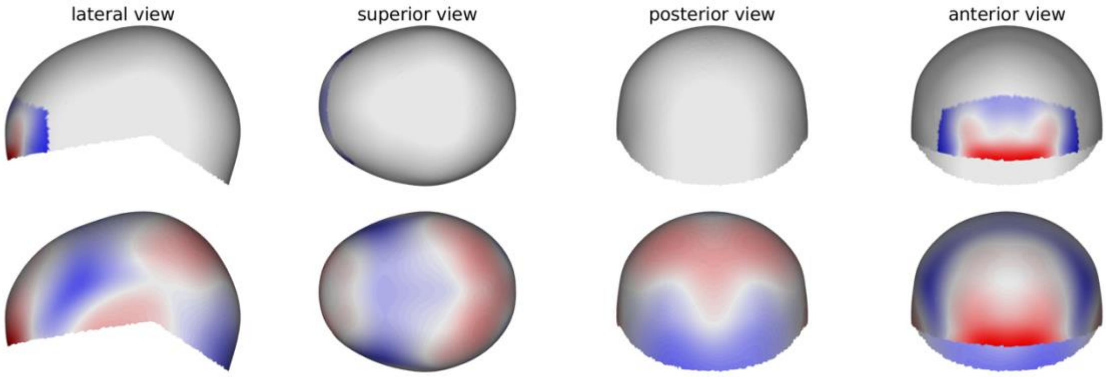

See page 2 for captions

**rs4842918**

**A**

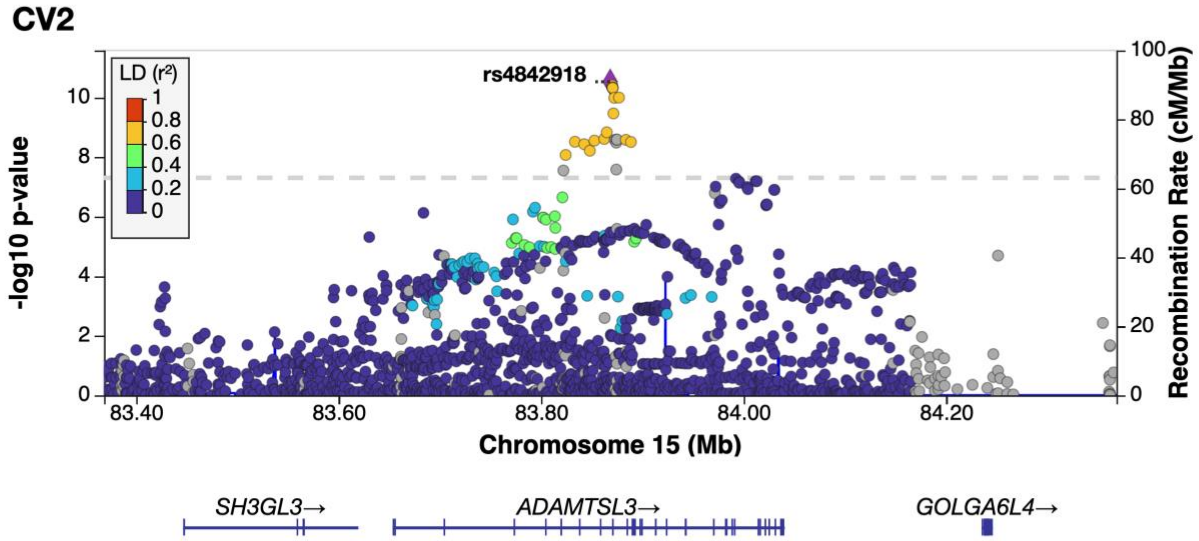

**B**

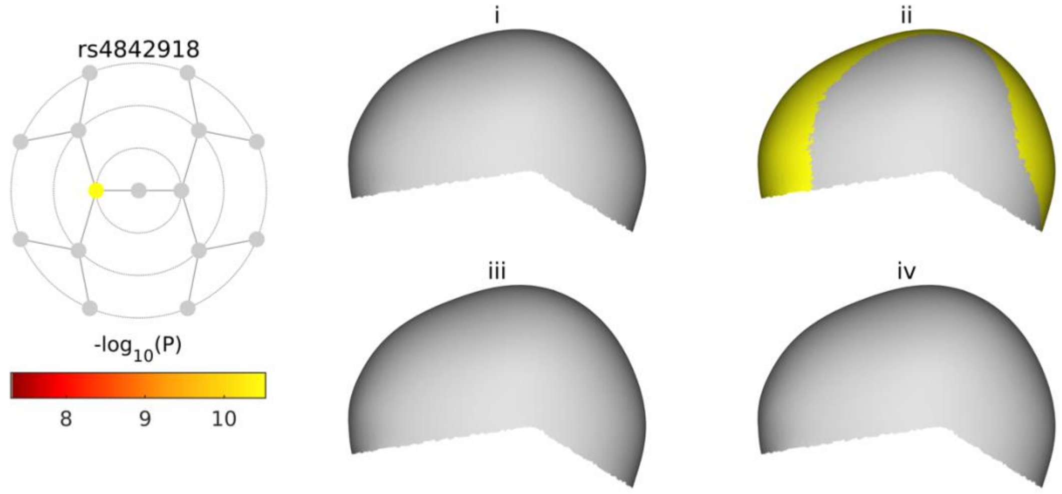

**C**

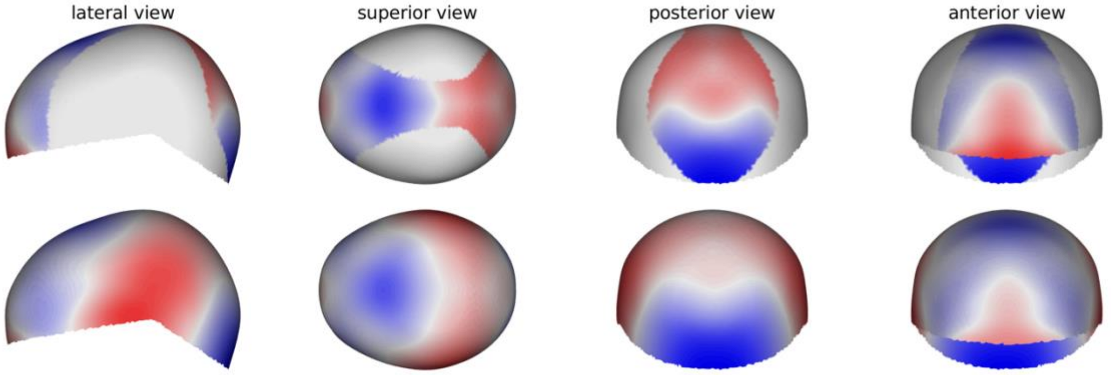

See page 2 for captions

**rs12940346**

**A**

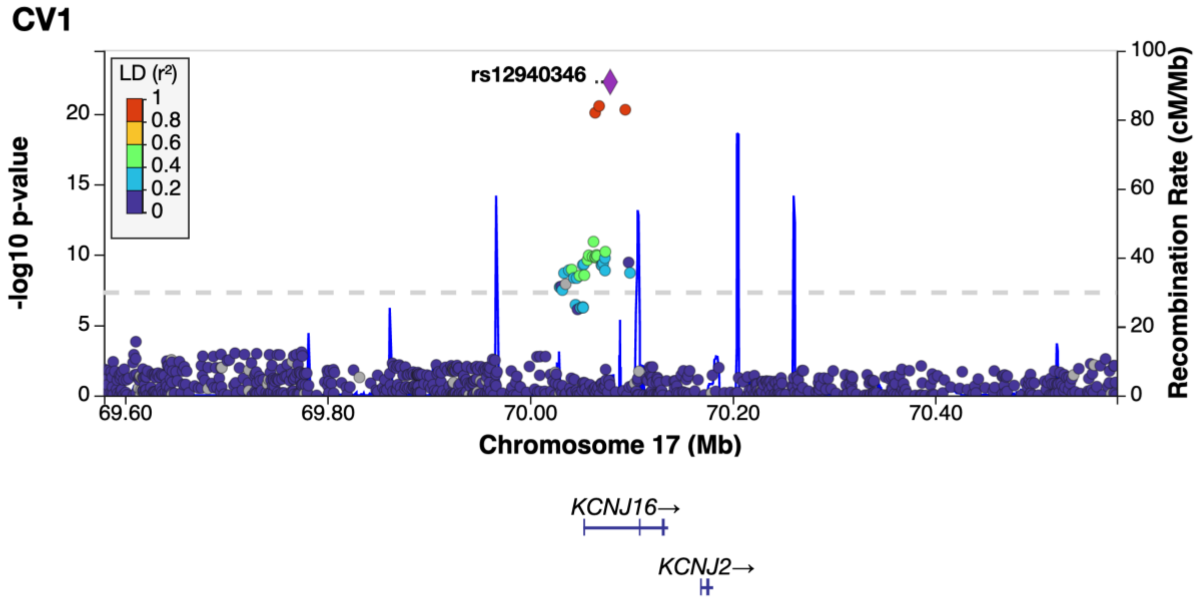

**B**

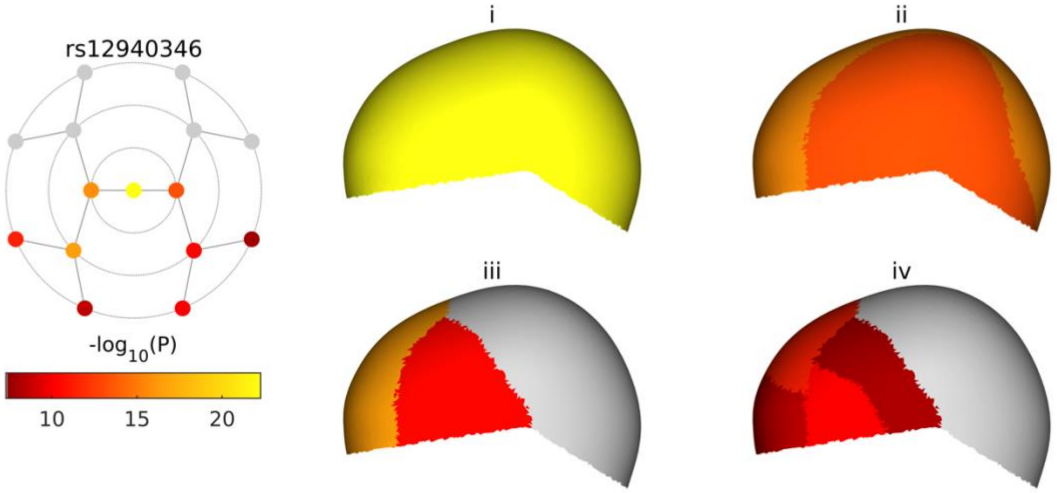

**C**

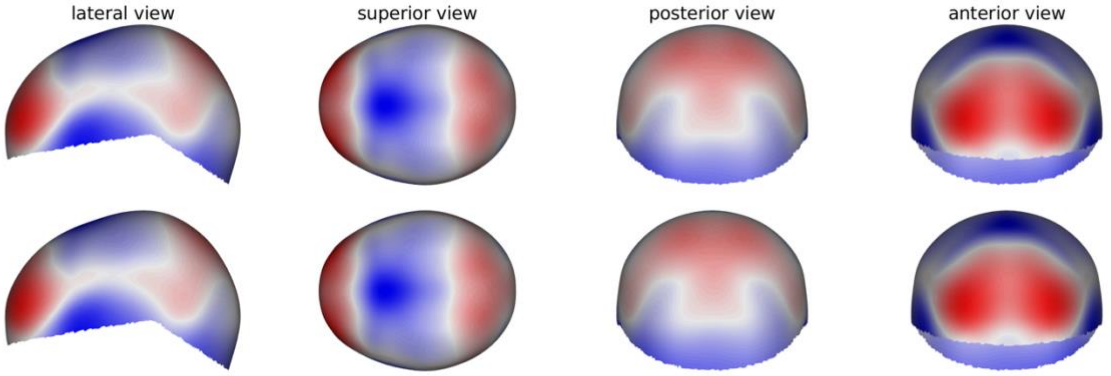

See page 2 for captions

**rs1321454**

**A**

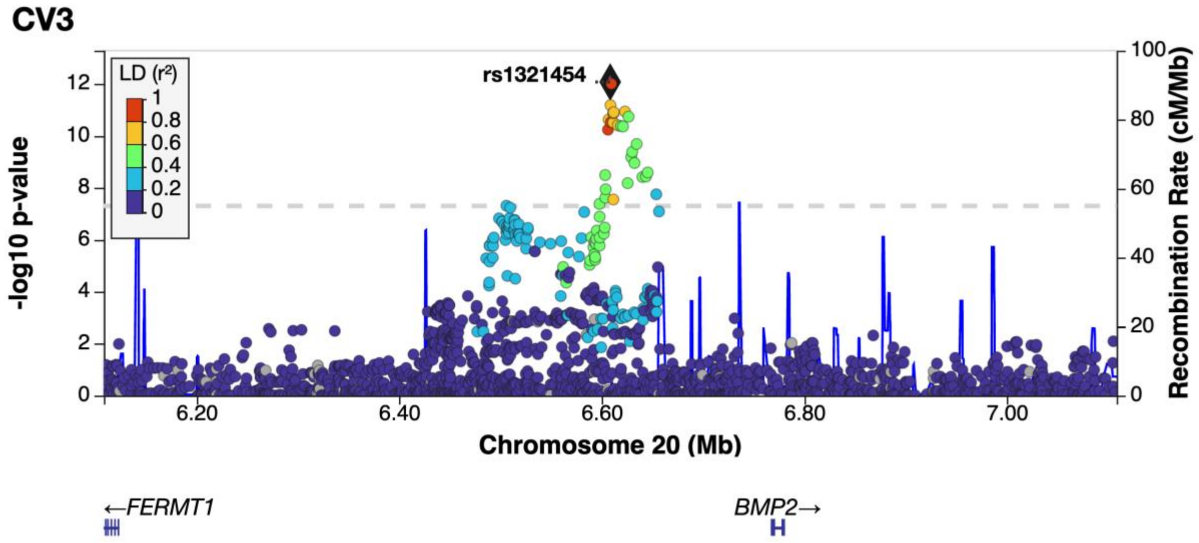

**B**

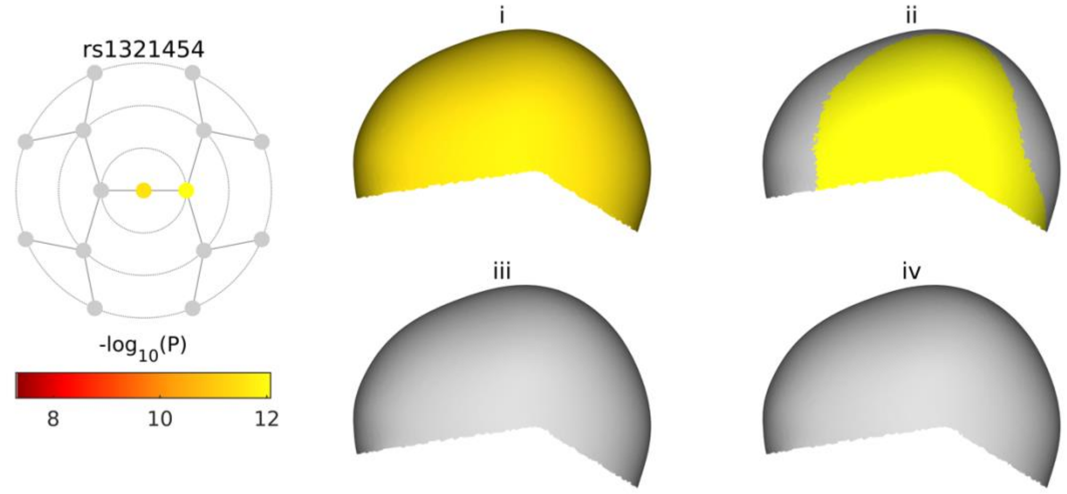

**C**

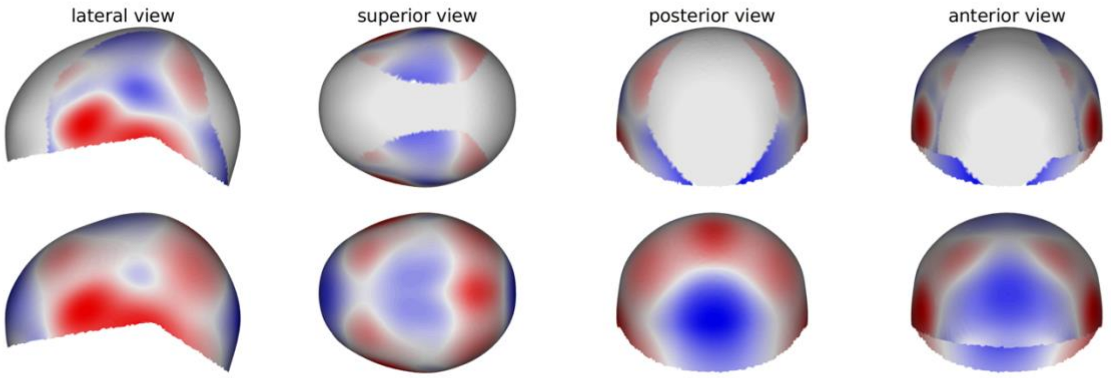

See page 2 for captions

**rs6054748**

**A**

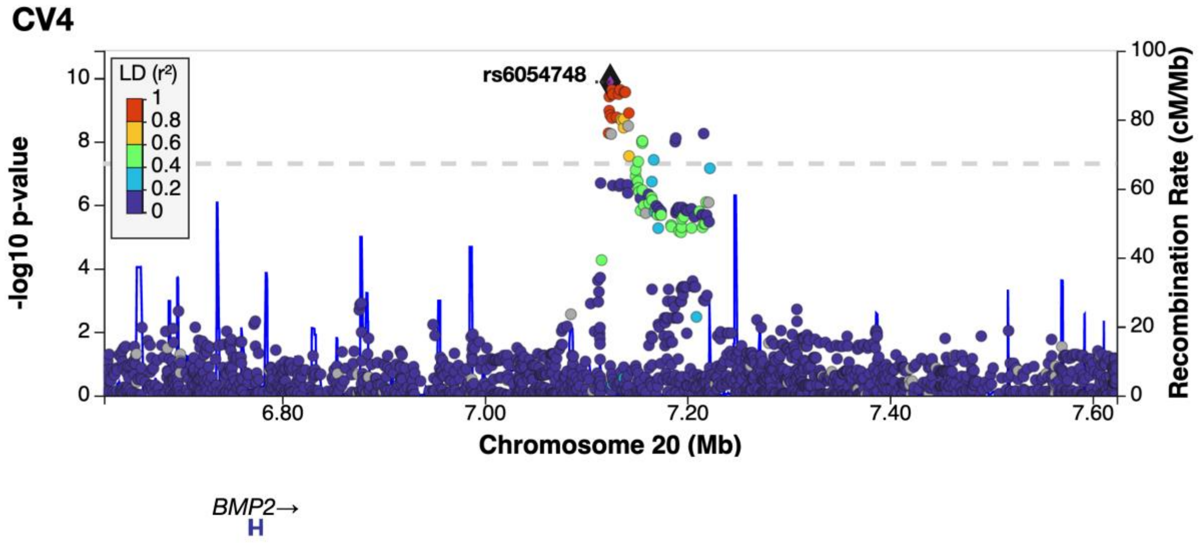

**B**

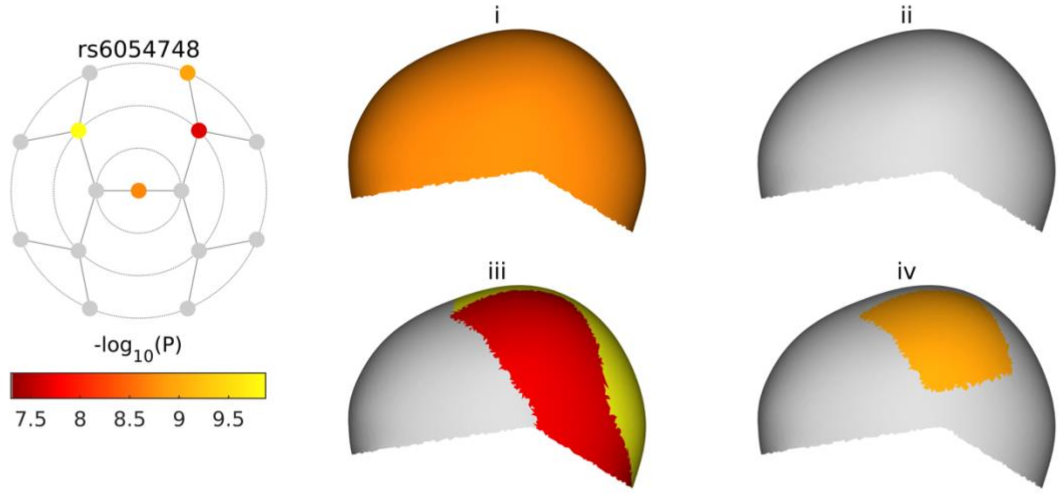

**C**

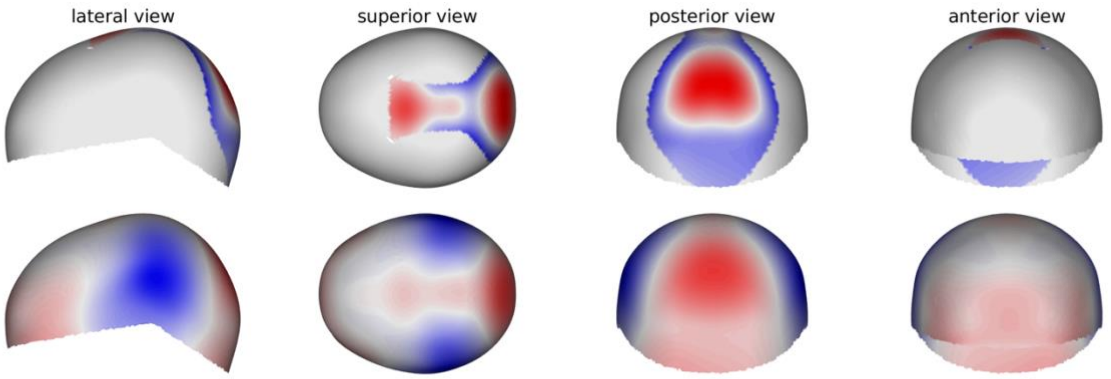
